# Norepinephrine Potentiates and Serotonin Depresses Visual Cortical Responses by Transforming Eligibility Traces

**DOI:** 10.1101/2021.06.22.449441

**Authors:** Su Z. Hong, Lukas Mesik, Cooper D. Grossman, Jeremiah Y. Cohen, Boram Lee, Hey-Kyoung Lee, Johannes W. Hell, Alfredo Kirkwood

## Abstract

Reinforcement allows organisms to learn which stimuli predict subsequent biological relevance. Hebbian mechanisms of synaptic plasticity are insufficient to account for reinforced learning because neuromodulators signaling biological relevance are delayed with respect to the neural activity associated with the stimulus. A theoretical solution is the concept of eligibility traces (eTraces), silent synaptic processes elicited by activity which upon arrival of a neuromodulator are converted into a lasting change in synaptic strength. Previously we demonstrated in visual cortical slices the Hebbian induction of eTraces and their conversion into LTP and LTD by the retroactive action of norepinephrine and serotonin Here we show *in vivo* in V1 that the induction of eTraces and their conversion to LTP/D by norepinephrine and serotonin respectively potentiates and depresses visual responses. We also show that the integrity of this process is crucial for ocular dominance plasticity, a canonical model of experience-dependent plasticity.

## Introduction

A fundamental role of our brain is to learn stimuli predicting biologically relevant values, such as novelty, saliency and reward. Neuromodulator signals have been thought to encode these value signals and teach the system which input should be reinforced ^1,2^ However, the evidence of biological value is often only available with temporal delay, which raises a question known as credit assignment problem^3,4^: How does the delayed value signal gate the plasticity in synapses that were transiently activated by the predictive stimulus? As a theoretical solution to bridge the temporal gap between stimulus and value signal, synaptic eligibility traces (eTraces) have been hypothesized. The eTrace is a transient and silent process, which is generated by pre- and postsynaptic activities of a neuron. The eTrace can last until the arrival of a neuromodulatory signal, forming a bridge between neuronal activities and behaviorally relevant neuromodulatory signals. This process allows selective plasticity of the synapses relevant to the biological values, while at the same time bridging the temporal delay that commonly exists between neuronal activity and feedback regarding its behavioral relevance^5–9^.

Synaptic eTraces were experimentally demonstrated first in the mushroom body of insects^10^, and later in several in vitro preparations of the mammalian brain, including striatum^11,12^, hippocampus^13,14^, and neocortex^15^. Particularly, we have shown the existence of two distinct eligibility traces (eTraces), the LTP and LTD traces, in primary visual cortex (V1) slices^15^. Specifically, the pre and postsynaptic activities generates the LTP trace at the synapse, which is converted into functional synaptic LTP by norepinephrine via involvement of β2-adrenergic receptors (β2AR) in layer 2/3 V1 pyramidal neurons. On the other hand, the post and then presynaptic activities generates the LTD trace, which is converted into synaptic LTD by serotonin via 5HT_2c_ receptors (5HT2cR)^15^. Notably, in contrast to several demonstrations of the eTraces *in vitro,* in slice preparations, the evidence of neuronal plasticity mediated by the transformation of the eTraces in vivo is limited to a single study in the striatum^16^. In the present study, we investigated whether the transformation of the LTP or the LTD trace in V1 result in the functional visual cortical response plasticity *in vivo*.

## Results

Previously we demonstrated *in vitro* the Hebbian induction of synaptic eTraces and its conversion into LTP and LTD by adrenergic and serotonergic receptors, respectively. Here we investigated the functional relevance of eTraces *in vivo* in the mouse visual cortex. To that end we tested 1) whether the release of neuromodulators timed in a reinforced-like paradigm modifies visual responses and receptive field selectivity 2) whether Hebbian associative paradigms can induce eTraces, and 3) whether blocking the conversion of eTraces into LTP and LTD prevents ocular dominance plasticity.

### Potentiation and depression of visual cortical responses by the retroactive action of norepinephrine and serotonin

We first tested whether the timely release of norepinephrine (NE) or serotonin (5HT) after visual stimulation potentiates or depress visual responses in V1 in a manner consistent with the transformation of eTraces. To that end we quantified visual cortical responses before and after conditioning with optogenetically-induced release of neuromodulators in mice expressing ChR2 in either noradrenergic (NE-ChR2) or serotonergic neurons (5HT-ChR2) (see Methods). Visual responses were elicited by drifting horizontal and vertical bars presented to one eye and recorded in the contralateral V1 using optical imaging of the intrinsic signal (ISI) (Fig. 1a). Conditioning consisted in the alternating presentation of fullfield horizontal and vertical drifting gratings, where the horizontal presentations, but not the vertical ones, were immediately followed by direct LED illumination of the exposed cortex (5s train of 470 nm pulses of 10 ms at 20 Hz; Fig. 1b).

**Figure 1.**
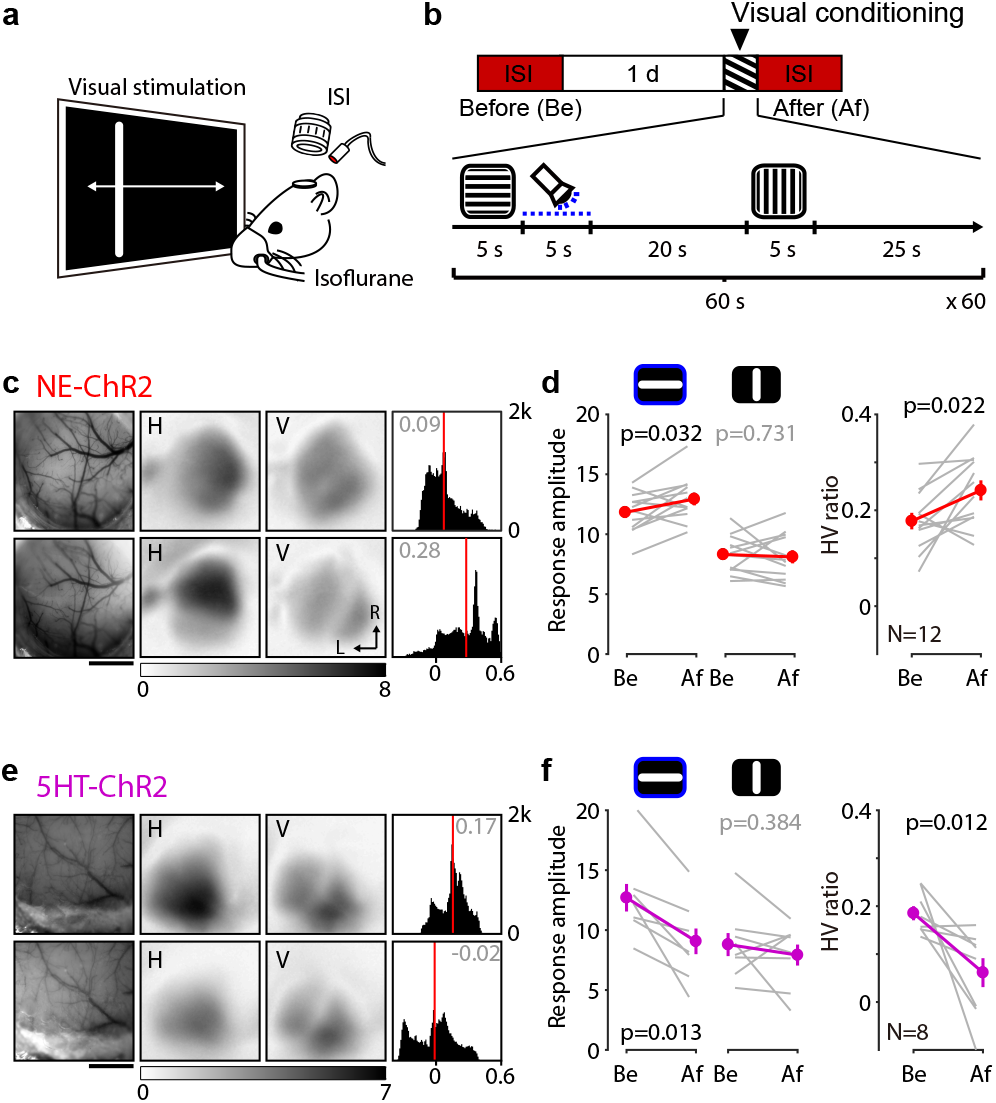
Potentiation and depression of visual cortical responses by the retroactive action of norepinephrine and serotonin. **a**, Schematic of the optical imaging of the intrinsic signal (ISI) of the visual cortical response from the V1. **b**, Experiment timeline (top) and the visual conditioning protocol (bottom). Blue dotted line indicates photoactivation of ChR2. **c-d**, Optogenetic transformation of LTP trace induces potentiation of the associated visual cortical response of the NE-ChR2 mice. **c**, Representative change of the visual cortical response by the visual conditioning. Left: vasculature pattern of the imaged region used for alignment. Scale bar, 1 mm. Middle: magnitude map of the visual cortical response evoked by horizontal [H] or vertical [V] drifting bar. Gray scale (bottom): response magnitude as fractional change in reflection x10^4^. Arrows: L, lateral, R, rostral. Right: histogram of HV ratio illustrated in the number of pixels (x-axis: HV ratio, y-axis: number of pixels). **d**, Summary of changes in response amplitude evoked by the horizontal (left) or vertical (middle) drifting bar as well as the change of HV ratio (right) before (Be) and after (Af) the conditioning. Thin line: individual experiments; thick line and symbols: average ± s.e.m. **e-f**, Optogenetic transformation of LTD trace induces depression of the associated visual cortical response of the 5HT-ChR2. Same format with (c and d).

In NE-ChR2 mice this conditioning resulted in substantial potentiation of the responses to the horizontal orientation, without changes in the responses to the unpaired vertical orientation (Fig. 1c,d). As a result, the ratio of horizontal versus vertical response amplitude (HV ratio), typically biased toward the horizontal orientation in naïve mice^17,18^, was further increased after conditioning. In contrast, in the 5HT-ChR2 mice the conditioning resulted in the selective depression of the horizontal responses without changes in the vertical responses and, consequently, a reduction in the HV ratio (Fig. 1e,f). Thus, the response to the same stimulation can be potentiated or depressed, depending on the neuromodulator released. As a control for the input specificity of the induced changes, in the NE-ChR2 we verified that the vertical responses are also modifiable by reinforcement-like conditioning (Extended Fig. 1a-d), and confirmed that conditioning the responses to the contralateral eye did not affect the responses evoked by the ipsilateral, non-conditioned, eye (Extended Fig. 1e-h). Altogether, the results are consistent with a scenario in which visual stimulation induces eTraces then can be converted to LTP and LTD by NE and 5HT respectively.

To test the involvement of eTraces more directly, we exploited our previous observation, made *in vitro,* that the conversion of eTraces into LTP and LTD by NE and 5HT requires the anchoring of β-adrenergic (βAR) and 5HT_2c_ serotonergic receptors to the postsynaptic protein PSD95^15^. Thus, peptides that mimic the C-terminal of these receptors disrupt their synaptic anchoring and prevent the conversion of the eTraces^15^. We tested, therefore, whether intraventricular infusionof cell-permeable versions of these peptides prior to the experiments prevent the modification of the visual responses by the reinforcement-like conditioning described above (Fig. 2a-d). In the NE-ChR2 mice injected with the peptide DSPL to disrupt the anchoring of βAR, the conditioning did not affect the HV ratio, whereas in mice injected with the control peptide DAPA^15^ the conditioning did increase the HV ratio (Fig. 2e). In a similar fashion, in 5HT-ChR2 mice injected with the peptide 2C-ct (TAT version of VNPSSVVSERISSV^15^) to disrupt 5HT2cR the conditioning failed to affect the HV ratio, yet the conditioning effectively reduced the HV ratio in control non-injected mice (Fig. 2f). We considered the possibility that prolonged exposure to the peptides might affect the regulation of intrinsic cell excitability by 5HT_2c_ and β-adrenergic receptors ^19^ and compromise the interpretation of the results. Arguing against that possibility, we found that in slices the application of the peptides did not affect the increase in layer 2/3 cell firing induced by 5HT_2c_ and ß-adrenergic agonists (Extended Fig. 2). This result also suggests that the conversion of eTraces, but not the regulation of cell excitability, is dependent on anchoring of these receptors to the PSD. Altogether then, the results are consistent with notion that the visual stimulation induced eTraces that were subsequently converted into LTP and LTD by the retroactive action of NE and 5HT, respectively.

**Figure 2.**
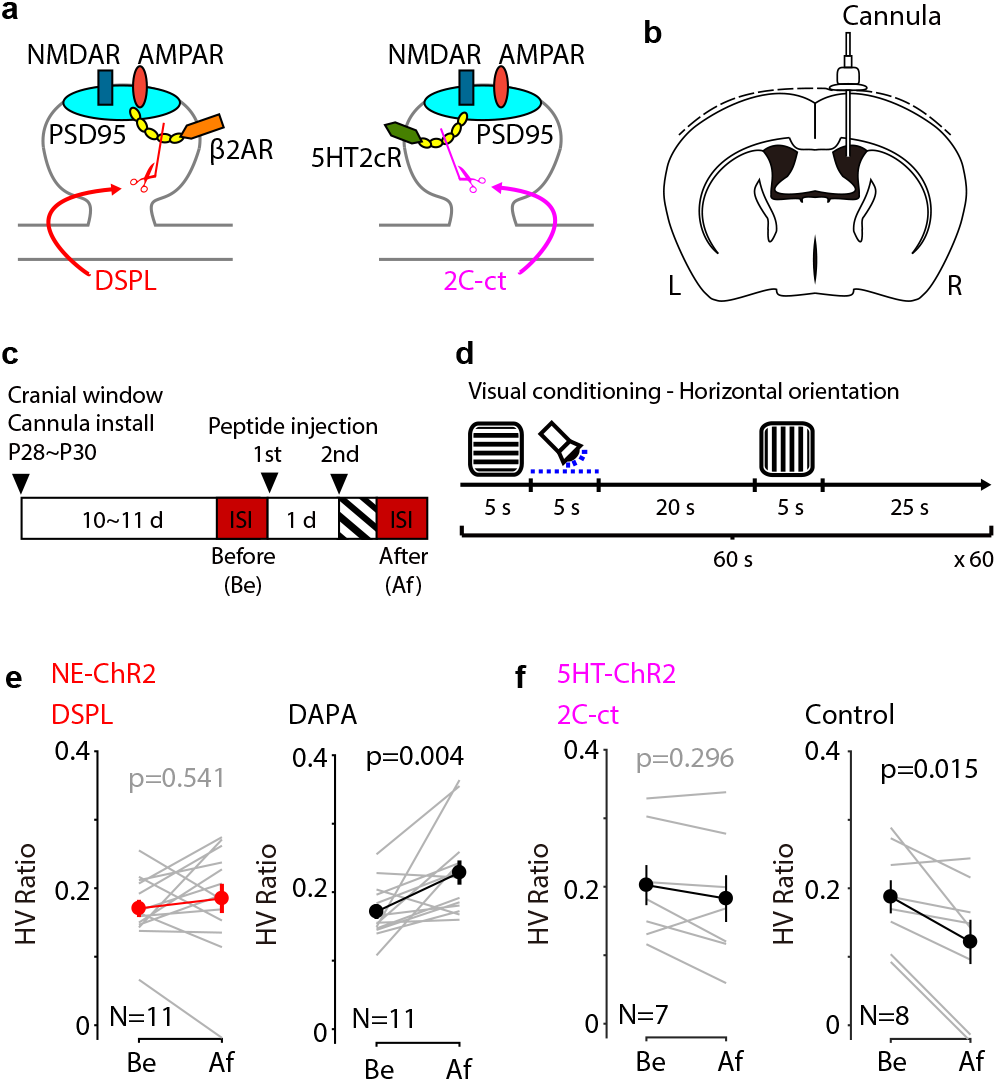
Peptides targeting the conversion of eTraces prevent reinforcement-like conditioning of visual cortical responses. **a**, Diagrams illustrating that DSPL (left) or 2C-ct (right) disrupts the direct interaction of the β2AR or 5HT2cR with PSD-95, respectively. **b**, A diagram illustrating the implantation of the cannula to inoculate the disrupting peptides to lateral ventricle. **c**, Experiment timeline. The peptide was injected 1 day (1st) and 30 min (2^nd^) earlier the visual conditioning. **d**, For the visual conditioning, horizontal or vertical drifting gratings were alternately shown with 30 seconds interval. Photoactivation to induce the release of neuromodulators was retroactively coupled to the horizontal drifting gratings. **e**, Summary of the HV ratio change by the visual conditioning in the presence of the disrupting peptide (DSPL) or the control peptide (DAPA) of the NE-ChR2 mice. **f**, Summary of the HV ratio change by the visual conditioning in the presence (2C-ct) or absence (control) of the 5HT2cR disrupting peptide of the 5HT-ChR2 mice. Thin line: individual animals; thick line and symbols: average ± s.e.m.

### Potentiation and depression of cortical visual responses by Hebbian induction and optogenetic conversion of eligibility traces

We previously demonstrated *in vitro* the induction of eTraces with Hebbian paradigms that associate pre-and postsynaptic activation. To examine that possibility *in vivo*, we tested whether pairing subthreshold visually evoked postsynaptic potentials (VEPSPs) with postsynaptic firing can result in LTP or LTD if followed by optogenetic delivery of endogenous monoamines in the NE-ChR2 and 5HT-ChR2 mice. We performed *in vivo* whole-cell recording from excitatory neurons in the superficial layers of V1 (Fig. 3 and Extended Fig. 3, see Methods) and subthreshold VEPSPs were evoked by two alternating rectangular flashing lights (500 ms) selected from 15 non-overlapping subregions of a screen (Fig. 3a). During associative Hebbian conditioning each VEPSPs was paired with a long burst of postsynaptic spikes (induced by current injection; 400 ms, 726.6±170.1 pA, 10.9±3.3 spikes, mean ± s.d.; Fig. 3d) that preceded and overlapped with the visual stimulation. The purpose of this temporal arrangement was to ensure post-pre and pre-post synaptic associations that we had shown *in vitro* to induce eTraces for LTP and LTD^15^. During the conditioning, optogenetic release of neuromodulators (1 sec train of 470 nm LED pulses, 10 ms at 20 Hz) was immediately coupled to one of the VEPSPs (C-VEPSP), the other VEPSP uncoupled to neuromodulator release (U-VEPSP) served as an internal control.

**Figure 3.**
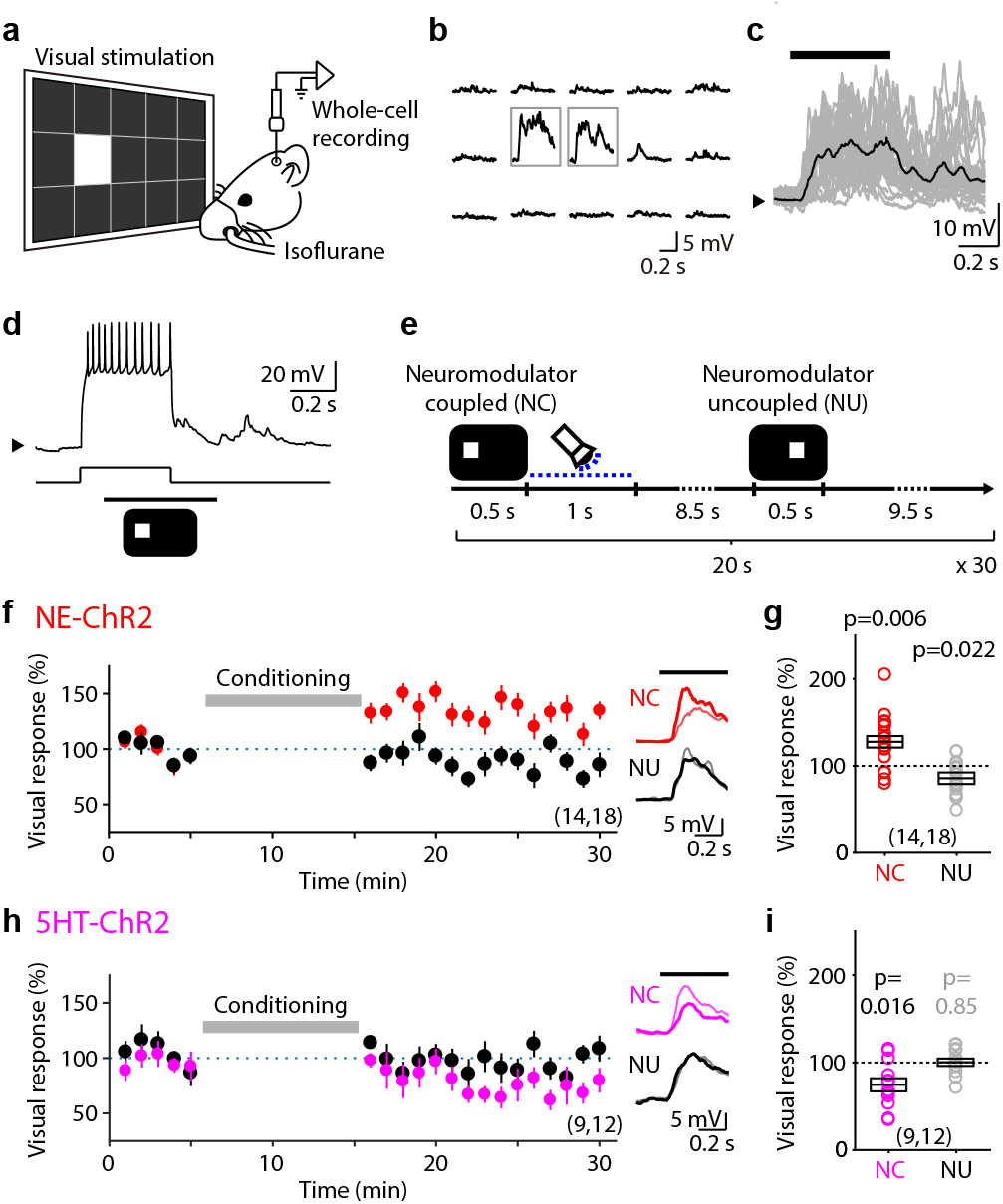
Potentiation and depression of VEPSPs by Hebbian induction and optogenetic conversion of eligibility traces. **a**, Schematic of *in vivo* whole-cell patch clamp recording of the superficial V1 neurons. **b**, Example of the VEPSPs elicited by visual stimuli at each subregion of a screen. Gray boxes indicate the two panels chosen for visual stimulation. **c**, Individual (gray traces) and averaged (black trace) VEPSPs elicited by the visual stimulus presentation (black bar on top) of a representative neuron. Black arrow on the left indicates Vm (−70 mV). **d**, Pairing of VEPSP with a burst of postsynaptic spikes. The current injected via recording pipette and the timing of visual stimulus is shown at bottom. Black arrow on the left indicates Vm (−70 mV). **e**, Visual conditioning protocol. **f**, Normalized change of the NC (red) or NU (black) VEPSP amplitude of the NE-ChR2 mice by the visual conditioning. Inset traces show the VEPSPs of a representative neuron averaged initial (thin line) or last (thick line) 5 minutes of the recording. **g**, Summary of VEPSP changes by the visual conditioning. Box plot: average ± s.e.m. Sample number indicates the number of animals and the number of recorded neurons. **h-i**, Same as panel (f) and (g), but for the 5HT-ChR2 mice.

To test the transformation of LTP traces, we recorded in NE-ChR2 mice (TH-ChR2, see Methods) and the optogenetic release of NE was evoked by direct illumination of the cranial window (Extended Fig. 3a,b, see Methods). In accord with our previous *in vitro* findings^15^ after the conditioning the C-VEPSPs were potentiated (p=0.006; Fig. 3f,g). The U-VEPSPs, on the other hand, were slightly, but significantly depressed (p=0.022) after the visual conditioning (Fig. 3f,g). These changes were activity dependent because the optogenetic activation alone did not affect the VEPSPs (104.4±7.8 %, p=0.635, n=13) (Extended Fig. 3f,g). In addition, the conditioning did not affect passive membrane properties of the cells (Extended Fig. 3d)

We examined the transformation of LTD traces by illuminating, via optic fiber, the dorsal raphe nucleus (DRN) of 5HT-ChR2 mice (Sert-Cre, Extended Fig. 3c, see Methods). Consistent with our previous slice results^15^, the conditioning depressed the C-VEPSPs (NC: 25.8±7.9%, p=0.016, n=12) without affecting the U-VEPSPs (NU: 0.3±4.4%, p=0.85, n=12) (Fig. 3h,i). Together, the results in the 5HT-ChR2 and NE-ChR2 mice demonstrate *in vivo* that the conjunction of synaptic activation and postsynaptic firing is sufficient to induce eTraces that subsequently can be converted into potentiation or depression by the retroactive actions of NE and 5HT.

### Optogenetic transformation of eligibility traces changes orientation responses of individual V1 neurons

A common feature of V1 neurons is their orientation selectivity, which is highly modifiable by sensory experience. We asked, therefore, whether reinforcement with optogenetic delivery of NE and 5HT can affect orientation responses of individual V1 neurons. On day 1 we recorded responses of the cells to drifting gratings of 6 orientations (in two directions) with two-photon calcium imaging in mice virally infected to express GCamp6f (see Fig. 4a,b and Extended Fig. 4). On day 2 the conditioning took place and the orientation response were determined again. During conditioning one orientation (which varied from mouse to mouse. See methods) was paired with optogenetic release of neuromodulators in a similar fashion as it was done with the Hebbian studies (Fig. 4b). The orthogonal orientation, which was not paired, was also delivered in an alternated fashion and served as a control.

**Figure 4.**
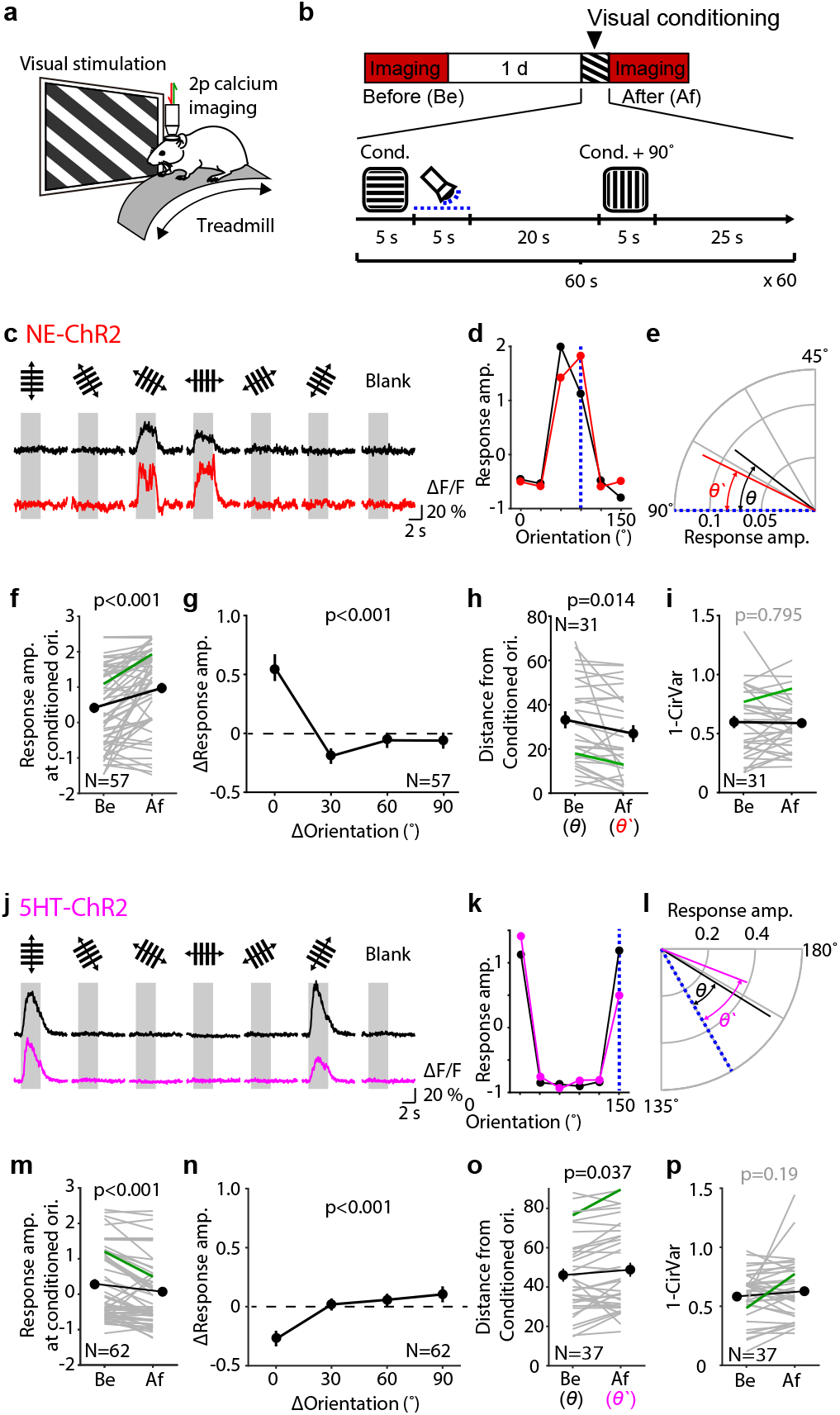
Optogenetic transformation of eligibility traces changes orientation responses of individual V1 neurons. **a**, Schematic of the two-photon calcium imaging of head-fixed mice. **b**, Experiment timeline (top) and the visual conditioning protocol (bottom). Blue dotted line indicates photoactivation of ChR2. **c-e**, Analysis of a representative neuron from a NE-ChR2 mouse. **c**, Fluorescence signal elicited by various orientations of the drifting gratings before (black) and after (red) the visual conditioning. Each trace indicates the average response across 16 trials consist of both directions. Gray area indicates the visual stimulus. **d**, Orientation tuning curve before (black) and after (red) the visual conditioning. Blue dotted line indicates the conditioned orientation. **e**, Vectors indicating the preferred orientation before (black) and after (red) the visual conditioning. Blue dotted line indicates the conditioned orientation. *θ* and *θ’* indicate the angular differences between the preferred orientations and the conditioned orientation. **f-i**, Summary of the changes in NE-ChR2 mice. **f**, Change of the response amplitude at the conditioned orientation. **g**, Response amplitude change according to the difference from the conditioned orientation. Data at two orientations (clockwise and counterclockwise) were pulled for the comparison. (Kruskal-Wallis test, KW stat = 42.58, p<0.001, and *post hoc* Dunn’s multiple comparison test (vs. 0° ΔOrientation, 30°: p<0.001, 60°: p=0.005, 90°: p=0.001). **h**, Change of the angular difference between the preferred orientations and the conditioned orientation after the visual conditioning. **i**, Change of the orientation selectivity by the visual conditioning. Thin line: individual neurons; thick line and symbols: average ± s.e.m. Green lines indicate the example neuron in (c-e). **j-p**, Summary of the changes in 5HT-ChR2 mice. Same format with (f-i). (Kruskal-Wallis test, KW stat = 30.59, p<0.001, and *post hoc* Dunn’s multiple comparison test (vs. 0° ΔOrientation, 30°: p<0.001, 60°: p<0.001, 90°: p<0.001)

We examined the potentiation of orientation responses by a reinforcement-like paradigm in 57 responsive cells recorded from 11 head-fixed awake NE-ChR2 mice (Fig. 4c-i). An example cell is shown in fig 4c-e. Before conditioning the cell responded best to a 60° orientation, after condition the cell became most responsive to the 90° used in the pairing (Fig. 4c,d). This particular cell had a clear orientation preference, which shifted towards the reinforced orientation after the conditioning; a change that was quantified as the angular difference between the preferred and the conditioned orientation (Fig. 4e). In the total population of examined cells, on average we observed an increased response that was specific to the conditioned orientation (Fig. 4f,g). In the subset of clearly orientation selective cells, the preferred orientation shifted towards the conditioned one (Fig. 4h), without losing overall selectivity. Those changes were not observed in control experiments in which the LED illumination was omitted during the visual conditioning (z-core before: 0.09±0.22; after:0.09±0.21; 3 mice, 22 cells; p= 0.997).

In 5HT-ChR2 mice we examined the depression of individual cell responses by the reinforcement-like paradigm. Initial experiments, performed as in figure 4c-e in head-fixed awake mice, resulted in no changes in the response amplitude to the conditioned orientation (z-core before: 0.99±0.15; after:0.93±0.15; 3 mice, 32 cells; p= 0.515) even after fluoxetine injection to reduce 5HT uptake (3 mice, 21 cells; p= 0.252). Therefore, we switched to the anesthetized preparation because it worked for the experiments previously described in Figures 1 and 2. Under these conditions, the pairing with the optogenetic stimulation of the dorsal raphe nucleus did result in the selective depression of the paired orientation. An individual example is shown in Fig. 4j-l; the summary averages of 62 cells in 6 mice are shown in Fig. 4m-p. It is presently unclear why the reinforcement-like paradigm with the current timing parameters were effective only in the anesthetized, and not in awake 5HT-ChR2 mice. Nevertheless, the results indicate that, *in vivo,* response preferences of V1 cells can be shifted away from or towards to a targeted orientation by the retroactive action of 5HT and NE, respectively.

### Preventing the transformation of eTraces blocks ocular dominance plasticity

In Fig. 1–4 we showed that optogenetical released NE and 5HT after visual stimulation can modify cortical responses in a manner consistent with the induction and conversion of eTraces. Because these results were obtained in anesthetized or head-fixed mice, it was of interest to examine the role of eTraces in cortical plasticity occurring in non-constrained, freely behaving mice. We focused on ocular dominance plasticity (ODP) induced by monocular deprivation (MD), a canonical model of sensory-induced cortical modification that is well established in mice. In juvenile mice MD results in an initial rapid depression of cortical responsiveness to the deprived eye; whereas at later ages, in young adults, MD manifest as a delayed potentiation of the responses to the non-deprived eye^20–22^. These changes have been attributed to LTD and LTP mechanisms^23^ (but see Turrigiano and Nelson, 2004^24^). We asked, therefore, how the 2C-ct and the DSPL peptides, respectively targeting the synaptic anchoring of 5HT_2c_ and β-adrenergic receptors, affect ODP. The changes in cortical response to the two eyes were monitored with intrinsic signal imaging in the binocular zone of V1 contralateral to the deprived eye^25^.

First, we tested whether the MD-induced depression of the deprived eye in juvenile mice (p28-29) requires the transformation of LTD trace and continuously infused the 2C-ct peptide, or its CSSA control, in the lateral ventricle starting 1 day prior the eye-suture (Fig. 5a-c). As expected from previous studies^20,22,22^, in control mice infused with the control CSSA peptide, 3 days of MD selectively reduced the responses to the deprived-eye (contralateral) without affecting the responses to the non-deprived eye (ipsilateral) (Fig. 5d,f,g). In contrast, in mice infused with the 2C-ct peptide the brief MD induced negligible changes the responses to either eye (Fig. 5e-g). A two-way ANOVA (Peptide x MD) and *post hoc* comparisons confirmed the significance of the interaction between the peptides and MD in the contralateral deprived-eye response (F(1, 18)=15.2, p=0.001) as well as in the ocular dominance index (ODI: see methods) (F(1, 18)=22.63, p<0.001). Thus, the normal MD-induced shift in response balance towards the non-deprived eye, quantified as a reduction in ocular dominance index (ODI:see methods), failed to occur in CSSA-infused mice (Fig. 5h). These results support the idea that conversion of eTraces for LTD by 5HT_2c_ receptors is necessary for juvenile ODP.

**Figure 5.**
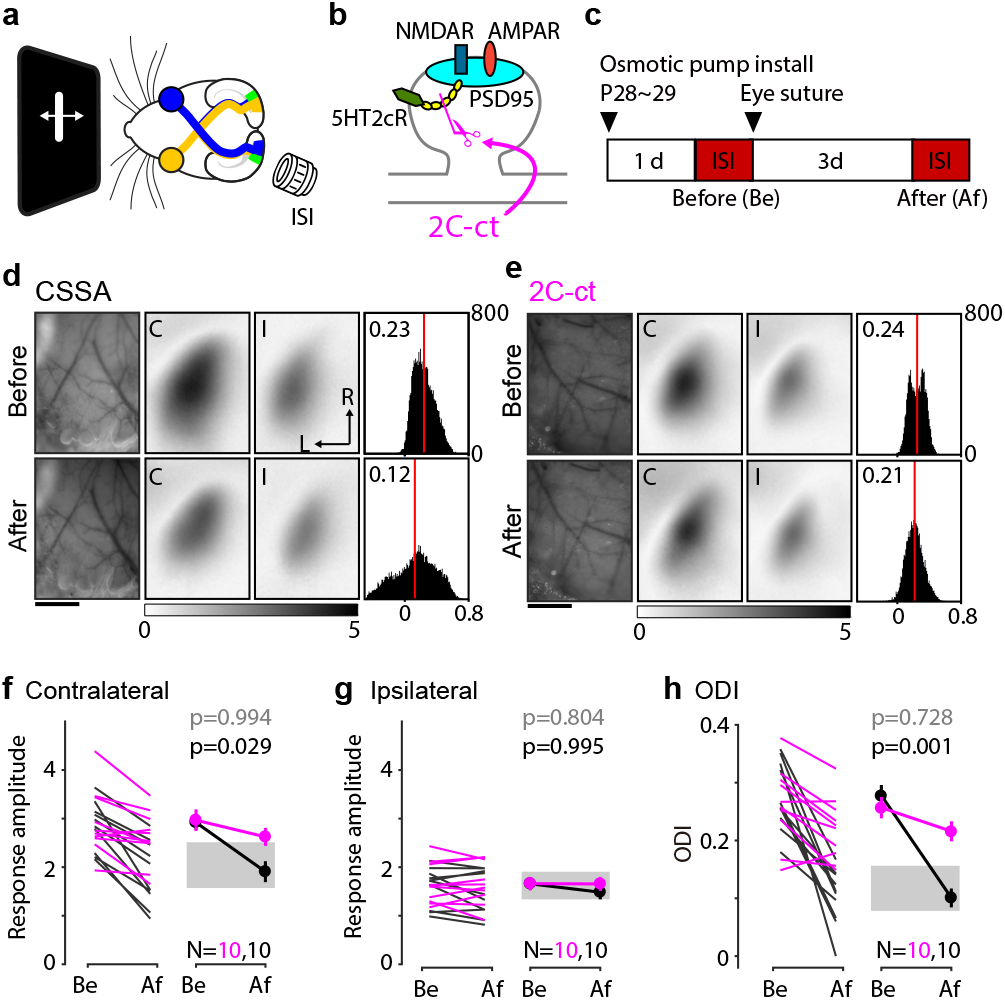
Blockage of the conversion of the LTD trace impairs the ocular dominance plasticity in juvenile mice. **a**, Schematic of the experiment to record the visual cortical response in the binocular region (green region at left hemisphere) of the V1. **b**, A diagram illustrating that 2C-ct disrupts the direct interaction of the 5HT_2c_ receptor with PSD-95. **c**, Experiment timeline. ISI were performed before (Be) and after (Af) the 3d MD. **d-e**, Representative change of the visual cortical response by the 3d MD in the presence of the control peptide, CSSA (d), or 2C-ct (e). Left: vasculature pattern of the imaged region used for alignment. Scale bar, 1 mm. Middle: magnitude map of the visual cortical response evoked by contralateral [C] or ipsilateral [I] eye from the recorded hemisphere. Gray scale (bottom): response amplitude as fractional change in reflection x10^4^. Arrows: L, lateral, R, rostral. Right: histogram of the ODI illustrated in the number of pixels (x-axis: ODI, y-axis: number of pixels). Red line indicates the average. **f-h**, Summary of the changes in response amplitude evoked by the contralateral (f) and ipsilateral (g) eye as well as the change of ODI (h) before (Be) and after (Af) the conditioning. Left: individual experiments in the presence of CSSA (black) or 2C-ct (purple); Right: average ± s.e.m. of left plot. p-values: Two-way ANOVA and *post hoc* Sidak’s multiple comparison’s test between the 2C-ct group and the CSSA group at before (gray) and after(black) the 3d MD. Gray region indicates 95% confidential interval of 3d MD mice without peptide infusion.

Next, we tested whether the transformation of the LTP trace is required for potentiating the non-deprived eye during prolonged MD (7 days) in young adult mice (p90-96) and evaluated the effects of infusing the DSPL peptide and the control DAPA peptide (Fig. 6). In these studies, infused and non-infused adult mice (p90-96) were subjected to MD for 7 days with the ocular dominance analyzed at the end of MD (Fig. 6a,b). As expected from previous studies, compared to the age matched normal reared mice, non-infused mice with 7d MD showed on average a potentiated response to the non-deprived eye. Consequently, their ODI was reduced, reflecting the shift toward the open eye (Fig. 6c,d). In contrast, mice infused with the DSPL peptide showed negligible potentiation of the nondeprived eye as well as minimally reduced ODI (Fig. 6g-i). On the other hand, mice with the control peptide DAPA did show the open-eye potentiation and reduced ODI comparable to the one registered in non-infused 7dMD mice (Fig. 6g-i). These results imply critical role of the transformation of LTP trace in the potentiation of open eye during the ODP.

**Figure 6.**
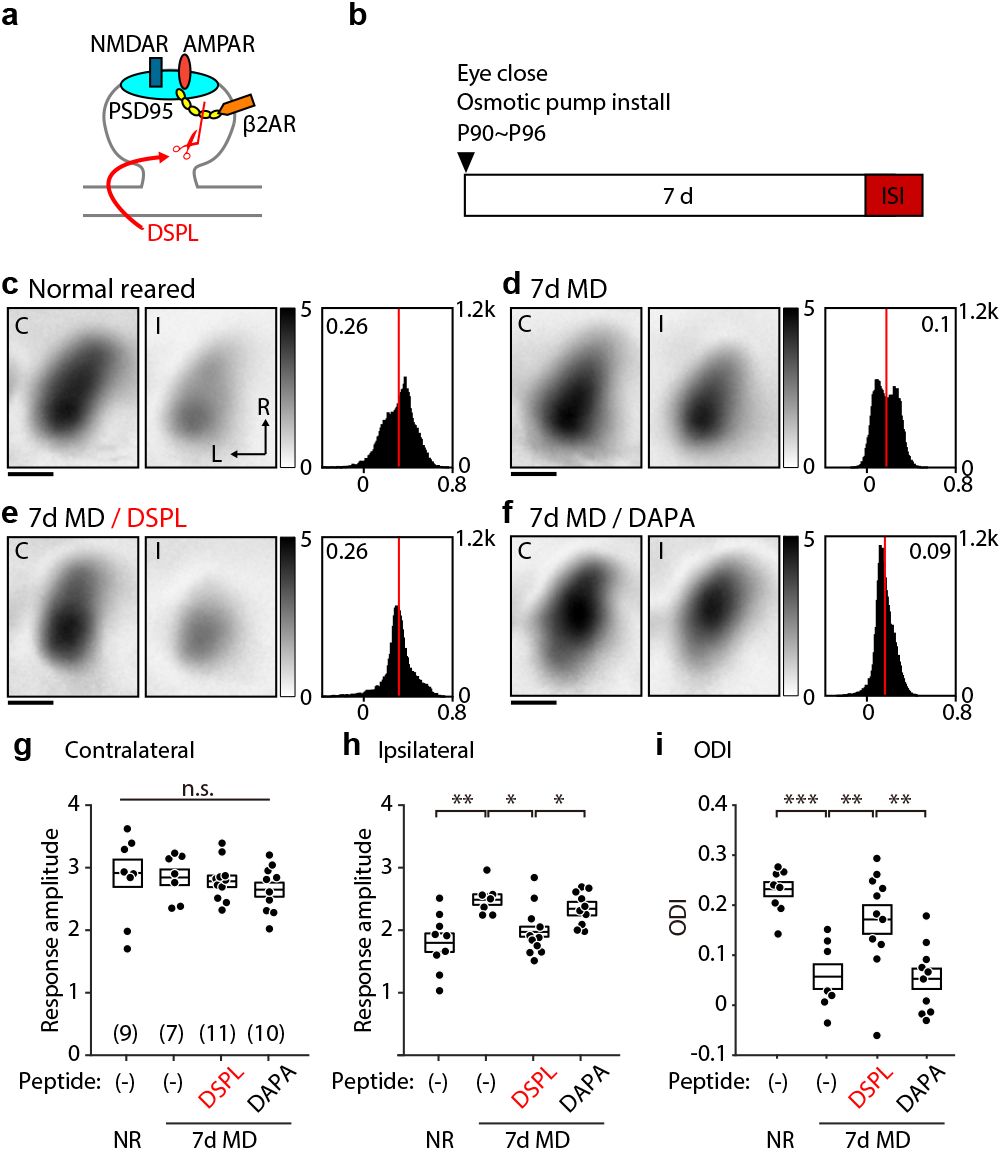
Blockage of the conversion of the LTP trace impairs the ocular dominance plasticity in young adult mice. **a**, A diagram illustrating that DSPL disrupts the direct interaction of the β2-adrenergic receptor with PSD-95. **b**, Experiment timeline. For the MD, contralateral eye to the recorded hemisphere was closed for 7 days before the ISI. Peptide infusion was started when the eye closed. **c-f**, Representative visual cortical response of the mouse normal reared (c), after 7 days of MD (d), after 7 days of MD with the infusion of DSPL (e), and after 7 days of MD with the infusion of the control peptide, DAPA (f). Left and middle: each magnitude map shows the visual cortical response from the contralateral (C) or the ipsilateral (I) eye from the recorded hemisphere. Gray scale: response amplitude as fractional change in reflection x10^4^. Arrows: L, lateral, R, rostral. Right: histogram of the ODI illustrated in the number of pixels (x-axis: ODI, y-axis: number of pixels). **g-i**, Summary of the response amplitude evoked by the contralateral (g) and ipsilateral (h) eye as well as the ODI (i) of each group. Box plot: average ± s.e.m. One-way ANOVA and *post hoc* Holm-Sidak’s multiple comparison test, (g) F(3,33) = 0.578, p=0.633; (h) F(3,32) = 6.267, p=0.001; (i) F(3,33) = 12.31, p<0.001; *p<0.05, **p<0.01, ***p<0.001.

Finally, to verify the specificity of the disrupting peptides in the interruption of the LTP or LTD trace transformation, we cross-examined the actions of the peptides and asked whether the 2C-ct affect the potentiation of open eye in adult mice, and whether DSPL impairs the depression of closed eye in juvenile mice. In adult mice infused with 2C-ct, the non-deprived eye potentiation induced by 7d MD was normal; and in juvenile mice infused with DSPL the depression of the deprived eye after 3d MD was also normal (Extended Fig. 5). This confirmation of the specific action of the peptides further validate the idea that the induction of eTraces and their conversion into LTP and LTD is a necessary process in ODP.

## Discussion

Our results provide direct evidence for a role of eTraces in experience-dependent cortical plasticity. We showed that the transformation of eTraces by the retroactive action of NE and 5HT is sufficient to potentiate and depress visual responses in V1 and showed that this process is necessary for ocular dominance plasticity. We also demonstrated the Hebbian induction of eTraces by pairing visual stimulation with postsynaptic firing. These findings line up neatly with the emerging notion of a “three-factor rule” for synaptic plasticity stating that besides STDP-like associations of pre-and postsynaptic activity, the expression of synaptic plasticity is also dependent on neuromodulator signaling of behavioral relevance or salience, including reward^2,9^.

The primary visual cortex is not merely an early stage of visual processing but also a primary locus of perceptual learning. Recently, evidence indicates that visual stimulation associated with delayed rewards can modify V1 function in mice and humans^26–31^. The general consensus is that neuromodulatory systems play a central role in this type of reinforced learning in primary sensory cortices by translating experience of reward and transmitting the respective signals^2,32,33^, yet the underlying mechanisms remain unclear. The induction and transformation of eTraces, which decay over seconds^15^, is an attractive and simple synaptic mechanism to enable the selective modification of visual responses associated with a delayed reward. Consider that typically in these visual perceptual learning tasks, thirsty subjects (rodents or humans) are rewarded with water a few seconds after visual stimulation, which results in stronger neural representations and better sensitivity to the conditioned stimuli^26–30^. These experimental settings and outcomes match well our finding that optogenetic delivery of NE immediately after an oriented stimulus selectively potentiates the cortical and cellular responses to that particular orientation (Fig. 1 and 3). It is also worth noting that the conversion of eTraces into LTP and LTD requires the anchoring of the β-AR and 5HT2cR to the synaptic PSD^15^, and the peptides we used to target that anchoring are quite specific at prevent that conversion without affecting the induction of LTP or LTD by other paradigms^15^, and without affecting the neuromodulation of intrinsic excitability (Extended Fig. 2). Based on these considerations we surmise that theses disruptive peptides could be used as a straightforward strategy to test the role of eTraces in the outcomes of perceptual learning tasks.

Most theoretical work and most of the experimental demonstrations of eTraces have focused on the case for LTP only. Complementary to the potentiation studies and consistent with a prior *in vitro* study^15^ demonstrating distinct traces for LTP and LTD, our results revealed that visual cortical activity generate eTraces that can be converted into LTP and LTD by NE and 5HT. Importantly, responses to the same stimulus were potentiated or depressed depending on whether it was paired with NE or 5HT. This indicates the visual activity generate both traces at the same time, and the results of single cell experiments (Fig. 3) indicates coexistence in the same cells. Whether both eTraces can coexist in the same synapses remains an intriguing question. Another unanswered question is the identity of the molecular mechanism underlying the induction and conversion of the eTraces. A plausible suggested scenario is that the LTP and LTD eTraces represent residual activity of kinases and phosphatases respectively and that ß-AR and 5HT2cR modify the phosphorylated state of the AMPAR promoting/enabling their trafficking in and out of the synapse^15^. Independent of the molecular mechanisms, the existence of distinct eTraces for LTD in addition to those for LTP has import consequences. Theoretical studies showed the presence of the two distinct eligibility traces enables learning reward timing and magnitude via the competition of the two traces^15,34^. It is also worth noting that the opposite/complementary function of 5HT and NE in visual cortical plasticity reported here, parallels the duality of actions proposed for 5HT and dopamine Daw et al, 2002^35^ and resonates with proposed roles for serotonin in cognitive flexibility^36,37^, supporting the generality of the two eTraces system as a mechanism of plasticity.

The essential role of neuromodulation in cortical plasticity is well stablished since the initial finding that ablation of noradrenergic function prevents ocular dominance plasticity^38^. Subsequent studies extended these observations to other neuromodulatory systems and other sensory cortices^39–44^. Initially neuromodulators were thought as permissive/enabling factors that promote the induction Hebbian synaptic plasticity via the enhanced cellular and network excitability associated with arousal and the awake state. It was later found that besides facilitating it, specific neuromodulators can also act directly at the level of the expression of plasticity to control its magnitude and polarity^45,46^. Aligned with this latter idea, disruptions of the serotonergic system or the noradrenergic system during ODP respectively prevent the closed-eye depression of the non-deprived eye response potentiation^47^. Our current findings point to an additional third level of control: retractive modifications in a reinforcement-like manner.

The cortical modifications induced by monocular deprivation are largely considered a form of unsupervised Hebbian learning in which cortical circuitry slowly become tuned to the statistic of the environment via the slow accumulation of small incremental synaptic changes. In this view, neuromodulation serves primarily to increase the gain of plasticity in a behavioral state dependent manner. In contrast, the induction and retroactive transformation of eTraces suggest a more punctuated pace of changes, with neuromodulators restricting the plastic changes only to moments of valuable biological experience. Our results with peptides that disrupt anchoring strongly support a role of reinforcement-like learning in ODP, although they do not rule out an independent contribution of unsupervised-like plasticity. Nevertheless, it must be noted that reinforced learning can be highly “efficient” ^2^. For example, 60 NE pairings within one hour induces a shift of magnitude comparable to that obtained after 7 days of monocular deprivation^21^. In addition, and in contrast to unsupervised learning, in reinforced learning only a small proportion of pre and postsynaptic coincidences evoked by vision result in synaptic plasticity. Considering that synaptic plasticity could be a metabolically demanding phenomenon^48–50^, restricting and subordinating visual cortical plasticity to behavioral relevance could be advantageous (i.e. Li et at, 2020)^51^. Finally, we would like to note that the demonstration of reinforcement-like learning via the transformation of eTraces in V1, somewhat unexpected in a primary sensory cortex, suggest the universality of the rules governing cortical plasticity.

## Competing Financial Interests

The authors have no competing financial interests

## Author Contributions

S.H, L.M and CG collected, analyzed and interpreted the imaging data, B.L and J.W.H made the DSPL and DAPA peptides, C.D.G and J.Y.C developed optogenetics in the 5HT-ChR2 mice, S.H, HKL and A.K. wrote the manuscript.

## Acknowledgements

We thank H. Shouval for valuable comments in the writing. Supported by grants R01DA042038 to JYC, R01 MH097887 and R01 AG055357 to JWH, R01-EY014882 to HKL, R01EY12124 and R01EY025922 to AK

---

**Extended Figure 1.**
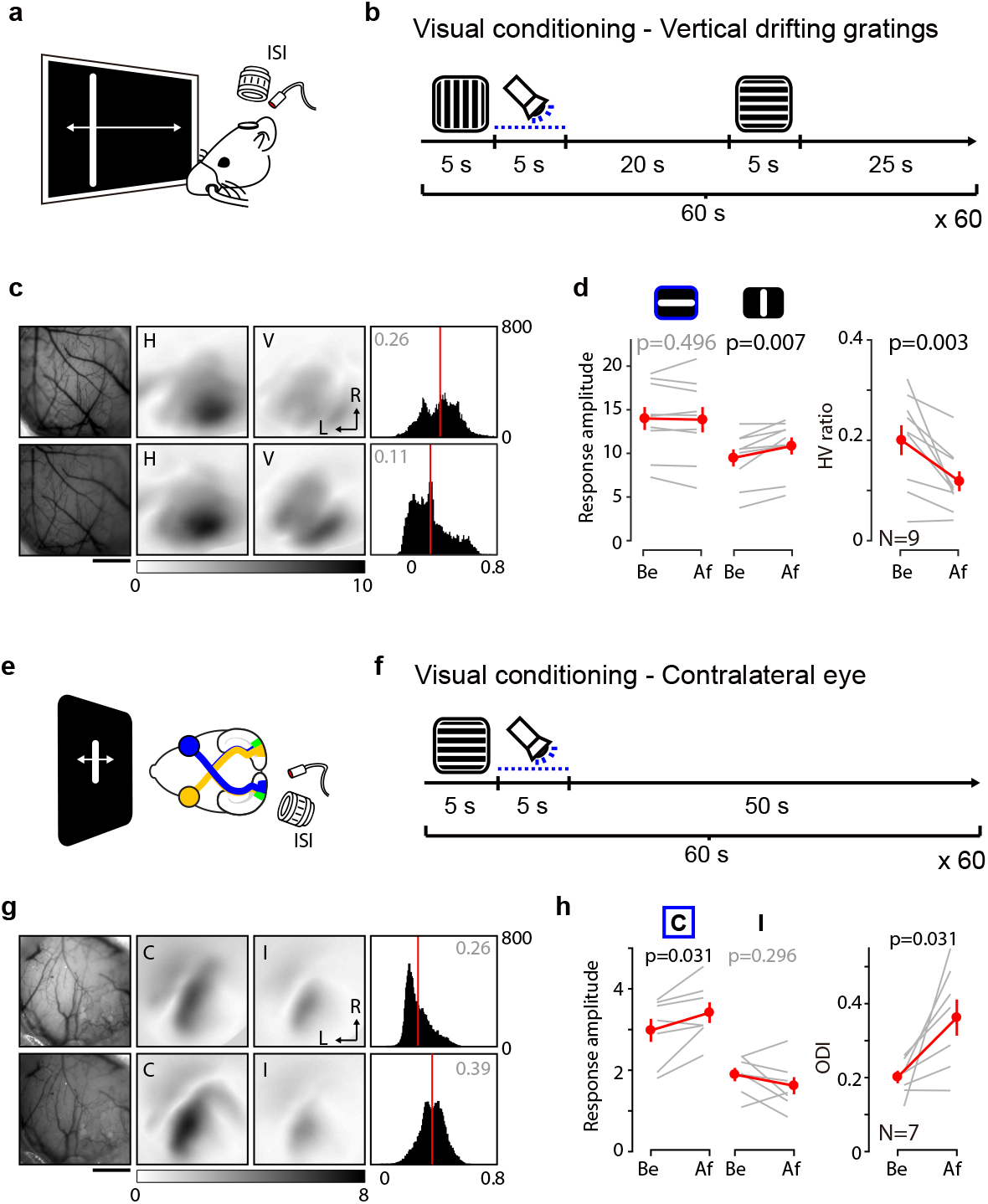
Further characterization of the potentiation of visual responses by optogenetical reinforcement in NE-ChR2 mice. **a-d**, Optogenetical reinforcement potentiates V1 responses to vertical drifting gratings. **a**,**b** Imaging of visual cortical responses was performed as in figure 1. **b**, Schematics of the conditioning protocol, similar to the one in figure 1, but with optogenetical stimulation paired to the vertical drifting gratings. **c**, Example responses before (top) and after (bottom) conditioning. Left: vasculature pattern of imaged region. Middle: responses to horizontal [H] or vertical [V] drifting bar. Right: histogram of the HV ratio illustrated in the number of pixels (x-axis: HV ratio, y-axis: number of pixels). **d**, Summary of the changes in response amplitude evoked by the horizontal (left) and vertical (middle) drifting bar as well as the change of HV ratio (right) before (Be) and after (Af) conditioning. **e-h**, Input specificity: conditioning one eye does not affect responses of the non-conditioned eye. **e**,**f** Experimental schematics. **e,** visual responses were imaged in the binocular region (green) of the V1. **f**, Conditioning protocol: the right eye was stimulated 60 times (1 per minute) with horizontal drifting gratings followed by the photoactivation to induce the release of norepinephrine. **g**, Example experiment. Left: vasculature pattern. Middle: visual cortical responses evoked by the eye contralateral [C] or ipsilateral [I] to the recorded hemisphere. Right: histogram of the ocular dominance index (ODI) illustrated in the number of pixels (x-axis: ODI, y-axis: number of pixels). **h**, Summary of the changes in response amplitude evoked by the contralateral (C, left) and ipsilateral (I, middle) eye as well as the change of ODI (right) before (Be) and after (Af) the conditioning. Gray scale at bottom of **c,g** represents the fractional change of reflection x10^4^Arrows in **c,g**: L, lateral, R, rostral. Scale bar in **c,g**: 1mm.Thin lines in **d,h**: individual experiments; thick lines and symbols: average ± s.e.m.

**Extended Figure 2.**
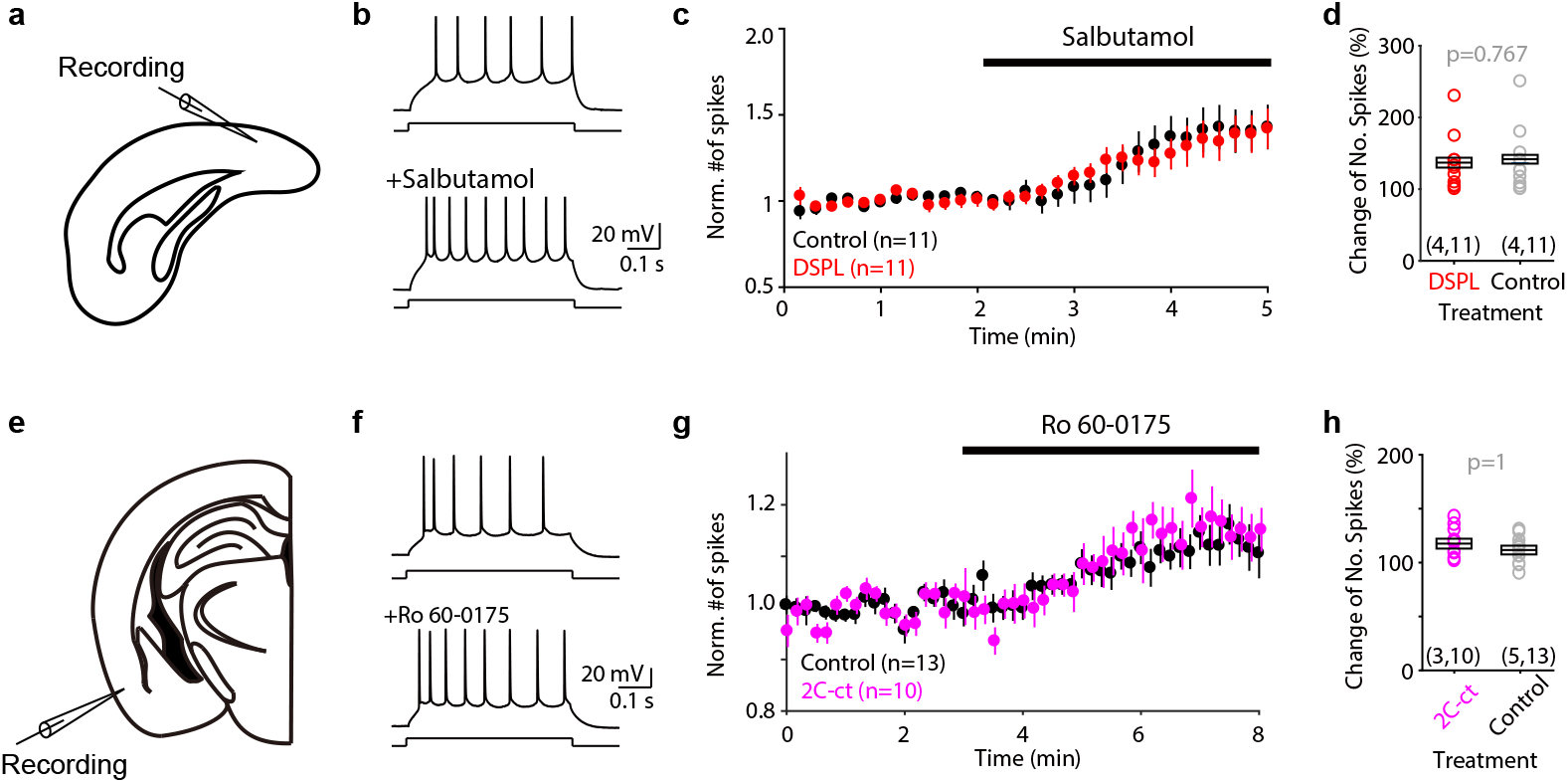
The DSPL and 2C-ct peptides do not affect the β2AR and 5HT2cR effects on intrinsic cellular excitability. **a**, Experiment diagram to test the excitability change by the β2AR agonist, salbutamol. The spike response of the layer 2/3 neurons in the V1 were recorded. **b**, Example spike response evoked by current step (bottom of the spike trace, 500 ms) before and after the administration of salbutamol. The current step amplitude was determined so that it evokes 5-6 spikes at the beginning of the recording and then maintained throughout the recording. **c-d**, Summary of the changes of the number of spikes evoked by the current step. The numbers of spikes are normalized by the average number of spikes during the baseline period (0-2 min). Slices were pre-incubated in the ACSF at least 20 min in the absence (black, Control) or presence (red) of DSPL. **e**, Experiment diagram to test the excitability change by the 5HT2cR agonist, Ro 60-0175. V1 neurons did not show excitability change by Ro 60-0175. Therefore, we tested the layer 2/3 neurons in the posterior piriform cortex, where the expression of the 5HT2cR is abundant (Clemett et. al., Neurophrmachology: 39, 123). **f**, Same in (b), but with Ro 60-0175. **g-h**, Same with (c,d), but with 2C-ct. The numbers of spikes are normalized by the average number of spikes during the baseline period (0-3 min). Box plot: average ± s.e.m.

**Extended Figure 3.**
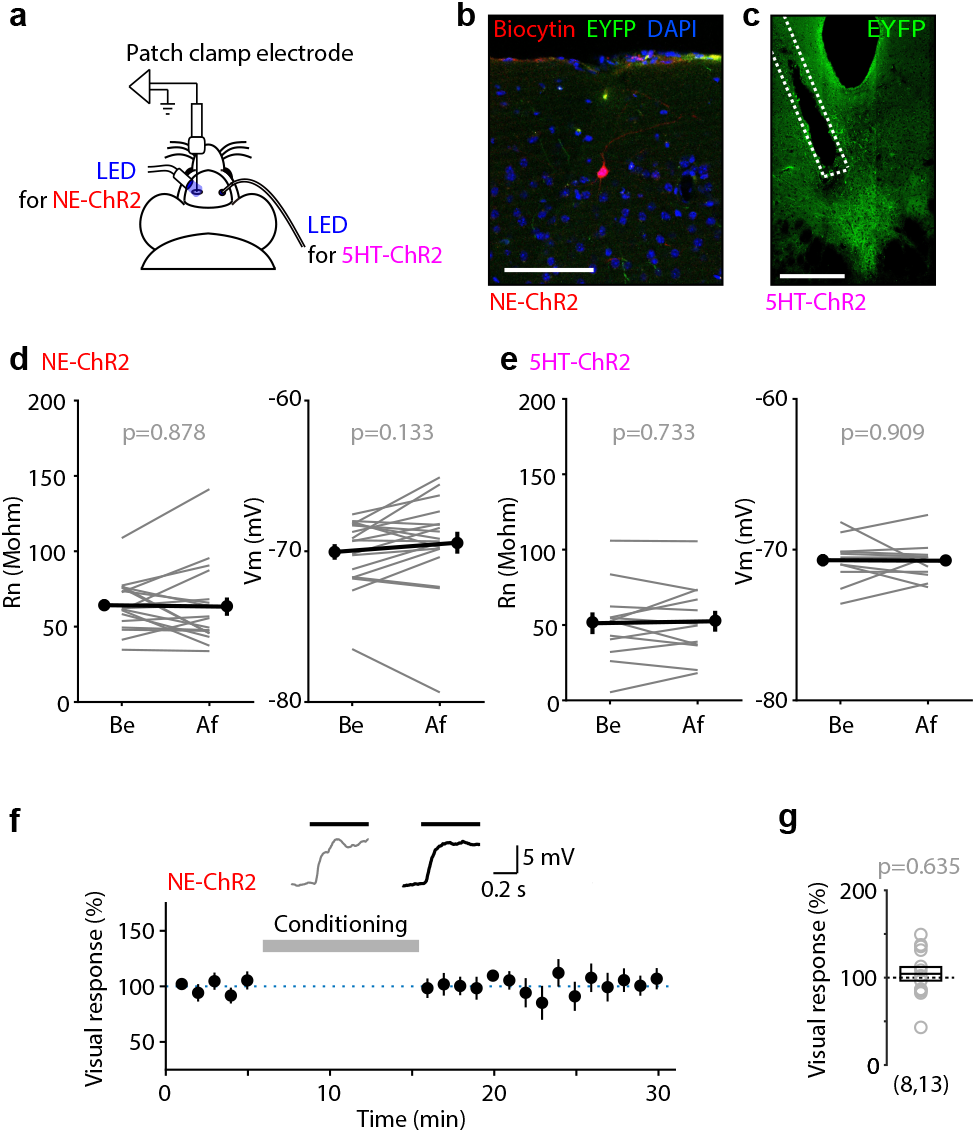
In vivo whole cell recording of VEPSPs. **a**, Photoactivation strategy schematics during the in vivo whole cell recording of the VEPSPs. Norepinephrine release was induced by photoactivation of adrenergic neuron terminal ChR2 of the NE-ChR2 mice at the V1 (left). Serotonin release was induced by photoactivation of the virally expressed ChR2 in serotonergic neurons via optic fiber implanted into the dorsal raphe nucleus (DRN) of the 5HT-ChR2 mice (right). **b**, An example fluorescence image from a NE-ChR2 mouse brain slice demonstrating the adrenergic neuronal projections expressing ChR2 (EYFP, green) and the recorded neuron (Biocytin, red). Nuclei are also visualized (DAPI, blue). Scale bar: 100 μm. **c**, An example fluorescence image from a 5HT-ChR2 mouse brain slice demonstrating the ChR2 expression (EYFP, green) in the serotonergic neurons at the DRN. White dotted line indicates the artifact by the implanted optic fiber. Scale bar: 100 μm. **d-e**, Input resistance (Rn) and base membrane potential (Vm) before and after the visual conditioning. **f-g,** Normalized change of NC-VEPSPs amplitude of the NE-ChR2 mice by the visual conditioning without pairing of the postsynaptic spikes (d) and the summary of change (e). Inset traces on top show the VEPSPs of a representative neuron averaged initial (gray) or last (black) 5 minutes of the recording. Box plot: average ± s.e.m.

**Extended Figure 4.**
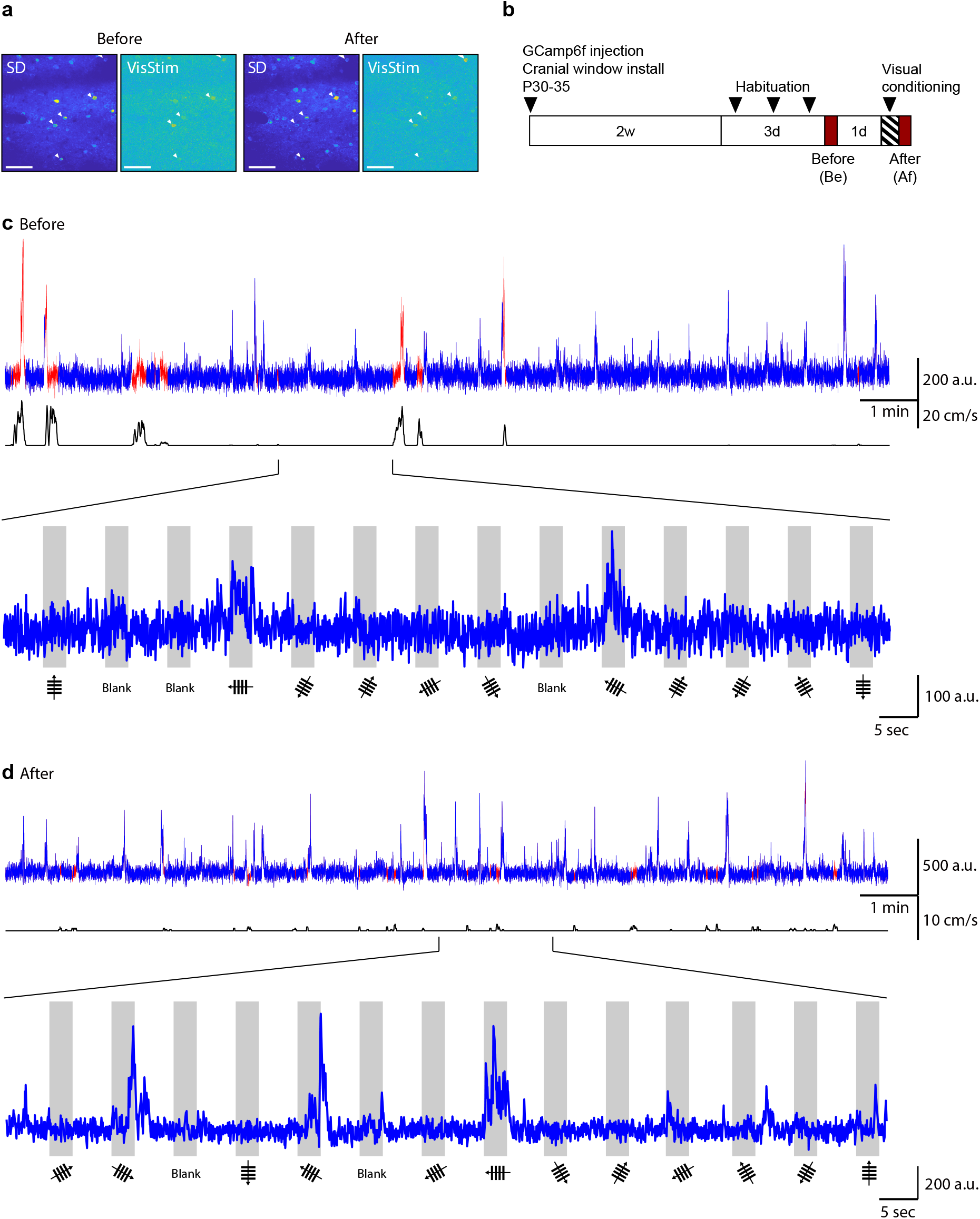
Two-photon calcium imaging of superficial layer excitatory neurons. **a**, Example two-photon imaging field of view visualized by fluorescence standard deviation (SD) or fluorescence correlation with the visual stimulus (VisStim) before and after the visual conditioning (see Methods). White arrows indicate the stable visually responsive neurons involved in the analysis. Scale bar: 100 μm. **b**, Experiment timeline. **c-d**, Top, fluorescence signal of the representative neuron across the recording time before (c) and after (d) the visual conditioning. Fluorescence signal during the stationary (blue) or during the locomotion (red) is illustrated based on the treadmill speed (black line). Bottom, a part of the fluorescence signal to 14 consecutive visual stimuli is demonstrated. Gray indicate the time the visual stimuli presented. The direction and the orientation of the visual stimuli are shown at the bottom.

**Extended Figure 5.**
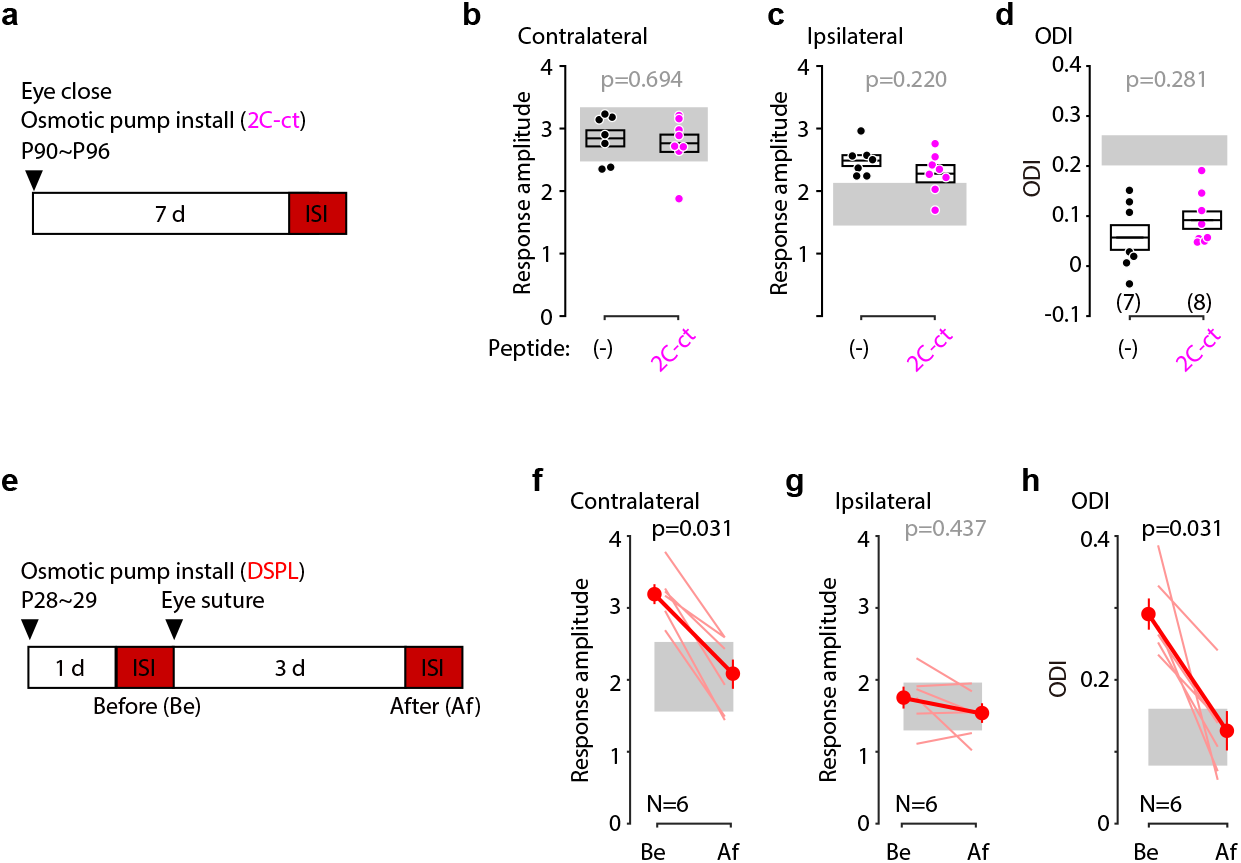
Specificity of the disrupting peptides in the interruption of the LTP or LTD trace transformation. **a-d**, Interruption of LTD trace transformation does not impair the potentiation of the open eye by the MD of young adult mice. **a**, Experiment timeline to test the specificity of the 2C-ct. **b-d**, Summary of the changes of the response amplitude evoked by the contralateral (b) and ipsilateral (c) eye as well as the ODI (d) of each experimental group. The data of 7d MD group were previously shown in Fig. 6. Gray region indicates 95% confidential interval values of normal reared mice. Box plot: average ± s.e.m. **e-h**, Interruption of LTP trace transformation does not impair the depression of the closed eye by the MD of juvenile mice. **e**, Experiment timeline to test the specificity of DSPL. **f-h**, Summary of the changes in response amplitude evoked by the contralateral (f) and ipsilateral (g) eye as well as the change of ODI (h) before (Be) and after (Af) the conditioning. Thin line: individual animals; thick line and symbols: average ± s.e.m. Gray areas describe the 95% confidence interval of the mice deprived for 3 days without the disrupting peptide.

## Online Methods

### Animals

All protocols were approved by the Institutional Animal Care and Use Committee (IACUC) at Johns Hopkins University and followed the guidelines established by the Animal Care Act and National Institutes of Health (NIH). In Figure 1–4 and Extended Figure 1–4, the mice at the age of P40-P80 were used. NE-ChR2 mice were produced by crossing THi-cre homozygote (provided by Dr. Jeremy Nathan) with Floxed-ChR2 (B6; 129S-Gt(ROSA)26Sortm32(CAG-COP4*H134R/EYFP)Hze/J (Jackson Laboratory, Bar Harbor, ME). For the 5HT-ChR2 mice, Tph2-ChR2 (B6;SJL-Tg(Tph2-COP4*H134R/EYFP)5Gfng/J) (Fig. 1–2 and 4, Extended Fig, 1–2 and 4) (Jackson Laboratory) or Sert-Cre mice (B6.129(Cg)-Slc6a4tm1(cre)Xz/J) (Fig. 3 and Extended Fig. 3) (Jackson Laboratory) were used. In Fig. 5–6 and Extended Fig. 5, C57BL/6J (Jackson Laboratory) mice at the age of P28-P30 (juvenile) or of P90-P96 (young adult) were used. Mice were reared in a 12 hours light/dark cycle.

### Slice electrophysiology

#### Preparation of cortical slices

Brain slices from mice (4-5 weeks) were prepared as described previously^1^. Briefly, mice were anesthetized using isoflurane vapors, then immediately decapitated. The brain was removed and immersed in the ice-cold dissection buffer (dissection buffer in mM: 212.7 sucrose, 5 KCl, 1.25 NaH_2_PO_4_, 10 MgCl_2_, 0.5 CaCl_2_, 26 NaHCO_3_, and 10 dextrose bubbled with 95%O2/5% CO2 (pH 7.4)). Thin (300 μm) coronal slices of visual cortex or piriform cortex were cut in the ice-cold dissection buffer and transferred to a light-tight holding chamber with artificial cerebrospinal fluid (ACSF in mM: 119 NaCl, 5 KCl, 1.25 NaH_2_PO_4_, 1 MgCl_2_, 2 CaCl_2_, 26 NaHCO_3_, and 10 dextrose bubbled with 95%O2/5% CO2 (pH 7.4)). The slices were incubated at 30°C for 30 minutes and then kept at room temperature until they were transferred to the disrupting peptide incubation chamber or to the recording chamber. For the disrupting peptide pre-incubation, the slices were incubated in the ACSF containing disrupting peptides (10 μM) at least for 15 min.

#### Whole-cell current clamp recordings

Recordings were made from layer 2/3 pyramidal neurons in normal ACSF with glass pipettes (3-5 Mohm) filled with potassium based internal solution (internal solution in mM: 130 K-gluconate, 10 KCl, 0.2 EGTA, 10 HEPES, 4 Mg-ATP, 0.5 Na-GTP, and 10 Na-phosphocreatine (pH 7.2-7.3, 280-290 mOsm). 500 ms square wave current step was injected to evoke 5-6 spikes every 10 sec and the number of spikes was monitored throughout recording. After 2-3 min of baseline recording, Salbutamol (40 μM) or Ro 60-0175 (10 μM) was perfused.

### Optical imaging of the intrinsic signal

#### Preparation of the NE-ChR2 mice

ISI of the NE-ChR2 mice were performed through a glass cranial window, which is prepared as described previously^2^ with some modifications. Briefly, 4-6 weeks old mice were anesthetized with isoflurane (2-3% for induction; 1.5% for maintenance in oxygen) and placed at stereotaxic frame. Dexamethasone (4.8 mg/kg, i.m.) and atropine (0.05 mg/kg, s.c.) were administrated to prevent brain edema and mucosal secretion, respectively. The skull was exposed and washed with hydrogen peroxide. The center coordinate of V1 was marked [−3.6/2.5] (A/P, M/L) and 3 mm diameter craniotomy was performed using dental drill. After removal of the bone flap, the three layered glass window^2^ was inserted and fixed with dental cement (C&B metabond, Parkell Inc.,NY). All procedure was performed under red LED and the glass cranial window was covered with non-transparent silicone sealant (Kwik cast; World Precision Instruments, Sarasota, FL) after the surgery to avoid the risk of activation of the terminal ChR2. The mouse was given an injection of Meloxicam (5 mg/kg, s.c.) and Carprofen (70 μg/ml) in the drinking water for 7 days following the surgery was given as analgesics. The mice were used for the experiment after at least 10 days of recovery period.

#### Preparation of the 5HT-ChR2 mice

ISI of the 5HT-ChR2 mice were performed through a glass window on the intact skull (Fig. 1) or a craniotomy (Fig. 2). To install a glass window on top of skull, the skull was exposed and washed with hydrogen peroxide and a 5 mm cover glass was mounted on top of the exposed skull with transparent cement (C&B metabond) [−3.6/2.5] (A/P, M/L). For the installation of the optic fiber (200 μm) (Thorlabs, Newton, NJ), a craniotomy was made at the coordinate of [−4.6/1.33] and the fiber was placed with 20° lateral angle targeting at the coordinate of [−4.6/0.2/-3.1] (A/P, M/L, D/V). Mice imaged through a glass cranial window was prepared as described above.

#### Preparation of the C57Bl/6 mice

Imaging window of the juvenile C57Bl/6 mice were prepared in the same way with 5HT-ChR2 above. For the preparation of imaging window of the young adult mice, the skull over the V1 region on the left hemisphere was exposed and washed with hydrogen peroxide. Agarose (3%) and a 5 mm round glass coverslip were placed on top of the exposed area.

#### Intracerebroventricular infusion of disrupting peptides

Cell membrane permeable LTP disrupting peptide DSPL (Myr-QGRNSNTNDSPL) and its control analogous peptide DAPA (Myr-QGRNSNTNDAPA) (gifts from J.W.H.) were prepared in 10% Dimethyl sulfoxide (DMSO) / 90% ACSF. LTD disrupting peptide 2C-Ct (TAT-VNPSSVVSERISSV) and its analogous control peptide CSSA (TAT-VNPSSVVSERISSA) (purchased from GenScript (Piscataway, NJ)) were prepared in ACSF. Each disrupting peptide was injected via cannula using syringe pump (2 ul, 660 uM) or using subcutaneously installed osmotic minipump (Alzet 1007D; Durect Corp., Cupertino, CA) combined with Brain Infusion Kit (Durect Corp.) (12 ul/day, 150 uM).

#### Acquisition and analysis of the intrinsic signal

ISI was performed following the method described previously^3,4^ with some modifications. Briefly, visual responses were acquired using a Dalsa 1M30 CCD camera (Dalsa, Waterloo, Canada). The surface vasculature and intrinsic signals were visualized with LED illumination (555-nm and 610-nm, respectively). The camera was focused 600 mm below from the surface of the skull. An additional red filter was interposed to the CCD camera and intrinsic signal images were acquired. The response amplitude of the intrinsic signal of an orientation and the HV ratio was calculated as following: (1) The cortical response at the stimulus frequency was extracted by Fourier analysis and the two maps generated with the opposite direction of drifting bar were smoothed by 5×5 low-pass Gaussian filter and averaged to generate the intensity map; (2) a combined intensity map was generated by sum of intensity map of the vertical and the horizontal orientation; (3) the region of interest (ROI) was defined by the region where the combined intensity is bigger than 40% of peak amplitude; (4) response amplitude of each orientation was computed by average of the intensity of all pixels in the ROI of each map; (5) HV ratio was calculated by the average of (H-V)/(H+V) of all pixels in the ROI, where H and V are the horizontal (H) and vertical (V) orientation, respectively. The response amplitude of the intrinsic signal from each eye and the ODI was also calculated by the same methods, but its ROI was defined at 30% of peak response amplitude of the smoothed intensity map from the ipsilateral eye, and the ODI was calculated by the average of (C-I)/(C+I) of all pixels in the ROI, where C and I are the contralateral (C) and ipsilateral (I) eye, respectively.

### In vivo whole-cell patch clamp recording

#### Preparation of the NE-ChR2 mice

For the preparation of cranial window on top of the V1, the mice were anesthetized with urethane (i.p., 1.2% in saline) and supplemented with isoflurane (0.5-1.2% in oxygen). Atropine was injected subcutaneously to reduce mucosal secretion (0.05 mg/kg) and eye drops were administered to keep eyes moist. The head skull was exposed and washed with 3% hydrogen peroxide. A head bar was attached to the anterior region of the head using dental cement (C&B metabond). The location of V1 was identified by stereotaxic coordinate (A/P: −3.6, M/L: 2.5). For initial trials, the location was confirmed using ISI. A small (~0.5 mm) cranial window was made with dental drill under red LED light and covered with 1% low melting point agarose (A9793, Sigma-Aldrich, St. Louis, MO) in the modified ASCF (in mM: 140 NaCl, 2.5 KCl, 11 Glucose, 20 HEPES, 2.5 CaCl2, 3 MgSO4, 1 NaH2PO4). Dura was not removed.

#### Preparation of the 5HT-ChR2 mice

5HT-ChR2 mice were produced by viral expression of Cre-dependent ChR2 in the dorsal raphe nucleus (DRN) of Sert-Cre mice. Specifically, craniotomy for the viral injection was made at the coordinate of [−5.46, 0] (A/P, M/L). Injection of AAVs (rAAV5-EF1a-DIO-hChR2(H134R)-EYFP) (Addgene, Watertown, MA) were performed with an angled approach (16°), in three different coordinates: [−4.66/-2.8], [−4.6/-3], [−4.54/-3.32] (A/P, D/V). For the installation of the optic fiber (200 μm) (Thorlabs, Newton, NJ), a craniotomy was made at the coordinate of [−4.6/1.33] (A/P, M/L). The optic fiber was placed with 20° lateral angle targeting at the coordinate of [−4.6/0.2/-3.1] (A/P, M/L, D/V). Mice were used for the recording experiment at least 2 weeks after the viral injection. Installation of a head bar and a cranial window for the recording was done as the same with the NE-ChR2 mice above.

#### Whole-cell current clamp recordings

Mouse body temperature was maintained at 37°C with heating pad and rectal probe and the heart rate was monitored throughout the experiment by electrocardiogram. Reference electrode was placed near the cranial window and submerged in the 1% agarose, which has been kept moist with the modified ACSF. Recordings were made with an Multiclamp 700B amplifier (Axon Instruments, Foster City, CA) using the blind patch-clamp technique^5^. Recording electrode (pipette resistance: 4-6 Mohm) with biocytin (1%) filled potassium based internal solution (in mM: 130 K-gluconate, 10 KCl, 0.2 EGTA, 10 HEPES, 4 Mg-ATP, 0.5 Na-GTP, and 10 Na-phosphocreatine (pH 7.2-7.3, 280-290 mOsm)) was used. Electrodes were inserted into the brain perpendicular and advanced with motorized micromanipulator (Sutter instrument, Novato, CA) in 1 μm increment. The depth of the recorded cell was estimated based on the depth from the pia and only the cells between 100 and 450 μm below the pia were used for analysis. Electrophysiological recordings and visual stimuli were controlled using acquisition software packages Stage (http://stage-vss.github.io) and Symphony (http://symphony-das.github.io). After the acquisition of whole-cell configuration, membrane potential was initially set to −70 mV by injection of hyperpolarizing current (30 – 260 pA) and was not adjusted throughout the recording. Input resistance (Ri) was monitored with hyperpolarizing current steps (50 pA, 100 ms) throughout the recording. The bridge was balanced, and liquid junction potential was not corrected. Sweeps were filtered at 2 kHz, sampled at 10 kHz and analyzed with custom code running in Matlab (The Mathworks, Natick, MA). Photoactivation to activate ChR2 was done via either the optic fiber placed in proximity of the recording cranial window (NE-ChR2) or the optic fiber implanted (5HT-ChR2). For the analysis of the VEPSPs, (1) voltage traces were smoothened by taking median over 50 ms window to eliminate the contamination by sporadic spikes during the visual response; and (2) the integrals of the voltage change during the flashing light illumination (500 ms) from the baseline (average voltage during 100 ms period prior to the onset of the flashing light) were calculated.

### Two-photon calcium imaging

#### Preparation of mice

The glass cranial window and the metal headpost to fixate the head during the recording was installed as described in Goldey et. al.^2^ For the expression of GCamp6f, 50 nl of AAV9-CamKII-Gcamp6f-WPRE-sv40 (Addgene) was injected 2-3 places near the central coordinate of the V1 [−3.6/2.5] (A/P, M/L), at a depth of 50 um. For 5HT-ChR2 mice, a craniotomy was made at the coordinate of [−4.6/1.33] (A/P, M/L) and the optic fiber (200 μm) (Thorlabs, Newton, NJ) was placed with 20° lateral angle targeting at the coordinate of [−4.6/0.2/-3.1] (A/P, M/L, D/V). Mice were used for the two-photon imaging at least after 2 weeks.

#### Two-photon calcium imaging and data analysis

Habituation of the mice to head fixation on a treadmill was done at least three times before the recording experiment. During the imaging session, the mouse freely moved on a treadmill and the locomotion was recorded using a quadrature encoder (US Digital, WA). Imaging was done on a custom built two-photon microscope (Janelia MIMMS) using Chameleon Ultra II laser (Coherent Inc., CA) and 8kHz resonant scanner (Sutter Instruments, CA). Images were acquired at ~30 fps using Scanimage 2018 (Vidrio)^6^ and analyzed using custom scripts written in Matlab. After image alignment, the region of interests (ROIs) were manually selected to analyze the visual stimulus responsive cells based on standard deviation and response to visual stimuli (Extended Fig. 4a). The response to visual stimuli was calculated at each pixel as below:

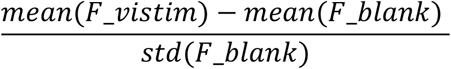

where F_vistim and F_blank indicate the fluorescence intensity by visual stimuli and blank stimuli, respectively, across the acquisition period. A semi-automated algorithm determined the shape of the template that was then used to extract fluorescent traces. Pixels on the boundary of these ROIs served as a local neuropil estimate. Fluorescence over time was measured by averaging within the ROI, and the fluorescence traces were subtracted by 0.7x neuropil estimate to correct for possible contamination by neuropil. The fluorescence traces during the mouse is moving (treadmill speed > 0.75 cm/sec) was not included in the analysis because it is known that the locomotion generates non-visual neuronal responses. ΔF/F was calculated as (F-F_0_)/F_0_, where F_0_ was the mean fluorescence intensity during a second prior to the onset of the drifting gratings. The relative response amplitudes by the drifting gratings at each orientation were demonstrated as z-scores after the average ΔF/F traces were calculated by taking mean (analysis with locomotion signal elimination for awake mice) or median (analysis without locomotion signal elimination for anesthetized mice) of 16 repeated traces. The orientation selectivity of neurons was determined by Kruskal-Wallis test and the preferred orientation of the orientation selective neurons (p<0.05) were determined as the orientation of a vector which is the summation of all vectors at each tested orientation.

### Visual stimulation and visual conditioning

All visual stimulation was presented on a LCD monitor screen diagonally placed 25 cm from the right (contralateral) eye of the mouse, except the visual stimulation for the recording of the ocular dominance where the screen was placed in front of the mouse.

#### Optical imaging of intrinsic signal to record the visual cortical response

Visual stimulation consists of a periodic vertical or horizontal drifting bar (2°) moving unidirectionally at 1 cycle/6 sec temporal frequency. Each visual stimulation was presented for 5 min. The order of the visual stimuli presentation was arranged such that the first and forth stimuli pair and second and third stimuli pair have the same orientations (i.e. direction order: (1) 270°, (2) 180°, (3) 0°, (4) 90°).

Visual conditioning consists of 30 repetition of visual stimulation blocks, in which square wave drifting gratings of 4 different directions (270°, 180°, 90°, 0°) with 1 cycles/20° spatial frequency and 1 cycle/6 sec temporal frequency are presented for 5 sec followed by 25 sec blank screen sequentially. The train of photoactivation (10 ms pulse at 20 Hz for 5 sec) followed the offset of the visual stimulus of the neuromodulator coupled orientation.

#### Optical imaging of intrinsic signal to record the ocular dominance at binocular region

Visual stimulation for the recording of the ocular dominance at the binocular region was restricted to the binocular visual field (−5° to +15° azimuth) and consisted of a horizontal thin bar (2°), continuously drifting for 5 minutes in upward (90°) and downward (270°) directions to each eye separately^1^. The sequence of the visual stimulation was arranged such that the first and fourth stimuli were for the same eye (270° and then 90°), and second and third for the other eye (90° and then 270°).

Visual conditioning consists of 60 repetition of visual stimulation blocks, in which square wave drifting gratings of 2 different directions (270°, 90°) are presented to the conditioned eye (right eye, contralateral to the recorded hemisphere) for 5 sec followed by 55 sec of a blank screen. The train of photoactivation (10 ms pulse at 20 Hz for 5 sec) followed the offset of the visual stimulus of the neuromodulator coupled orientation.

#### In vivo whole cell patch clamp recording

Visual stimulation consists of two non-overlapping rectangular flashing lights (500 ms) were alternately presented every 5 seconds. To decide the location of the stimuli, the screen was divided by 15 subregions and two of the subregions, which evoke reliable and comparable VEPSP, were selected at each cell. The contrasts of the stimuli were further adjusted so that their VEPSP amplitudes are comparable each other and remained subthreshold.

Visual conditioning consists of 30 repetitions of visual stimulation blocks, in which the two flashing lights at the selected subregion were alternately presented with 10 sec interstimulus intervals. For each visual stimulation, a square pulse current (400 ms) was injected via recording electrode 100 ms before the onset of the visual stimuli so that the evoked spikes and VEPSPs overlap significantly. The amplitude of current pulse determined at each cell to evoke at least 10 spikes by the current pulse. One of the two visual stimuli was chosen randomly (Neuromodulator Coupled, NC) and a train of photoactivation (10 ms pulse at 20 Hz for 1 sec) followed the offset of the NC visual stimulation. The other visual stimulus (Neuromodulator Uncoupled, NU) was alternately shown without photoactivation.

#### Two-photon calcium imaging

Visual stimulation consists of 16 repetitions of blocks, each of which containing square wave drifting gratings at 12 different orientations with 1 cycles/20° spatial frequency and 3 Hz temporal frequency and 2 blank stimuli. The stimuli were presented in a pseudorandom order, with each stimulus shown for 3 seconds followed by 5 seconds of blank screen.

For the visual conditioning, the neuromodulator coupled (NC) orientation was selected at each mouse based on the distribution of orientation preference of the recorded neurons in ‘Before session’ in a way that it is the orientation eliciting significant, but submaximal visual response. The visual conditioning consisted of 30 repetition of visual stimulation blocks, in which square wave drifting gratings of the NC orientation or the orthogonal orientation in two opposite directions are presented for 5 sec followed by 25 sec interstimulus intervals sequentially. The train of photoactivation (10 ms pulse at 20 Hz for 5 sec) followed the offset of the visual stimulus of the neuromodulator coupled orientation.

### Monocular deprivation

After the mice anesthetized with isoflurane (2-3% for induction; 1.5% for maintenance), the margins of upper and lower eye lids were trimmed and sutured shut. Small amount of Neosporin was applied to the sutured eye to prevent an infection. The mouse was given an injection of Meloxicam (5 mg/kg, s.c.) after the surgery. The sutured eye was checked before it was opened for an imaging session to make sure the integrity of the lid suture.

### Immunohistochemistry

The anesthetized mice were transcardially perfused with 10% neutral buffered formalin solution. Following perfusion, the brain was extracted and kept in the fixative solution overnight. The brain was sliced into 70 μm coronal sections and the slices were transferred to phosphate buffered saline (PBS). Slices were then permeabilized (2% Triton X-100 in 0.1 M PBS) for 1 hr before incubation with 1 mg/ml streptavidin-488 (in 0.1 M PBS containing 1% Triton X-100) overnight at 4°C. The slices were washed in PBS for an hour and mounted on a slide glass. Slides were coverslipped with the Prolong Gold anti-fade mounting solution with DAPI (Cell Signaling Technology, Inc., Danvers, MA) incorporated. Confocal images were taken on a Zeiss laser stimulated microscope 700.

### Statistical analysis

Normality was determined by Shapiro-Wilk test using Prism (GraphPad Software, San Diego, CA). Wilcoxon signed-rank test (Fig.1–3 and Extended Fig. 1,3,5), Wilcoxon ranksum test (Extended Fig. 2 and 5) or paired *t* test (Fig. 4) were performed using Matlab. Twoway ANOVA followed by *post hoc* Sidak’s multiple comparisons test (Fig. 5) and One-way ANOVA (Fig. 3 and 5) followed by *post hoc* Holm-Sidak test were performed using Prism. Data are presented as averages ± s.e.m. otherwise mentioned.

### Data Availability

Data are available from the corresponding author on reasonable request.

### Code Availability

Custom codes are available from the corresponding author on reasonable request.

## Notes

### Competing Interest Statement

The authors have declared no competing interest.

## References

1. Avery, M. C. & Krichmar, J. L. Neuromodulatory Systems and Their Interactions: A Review of Models, Theories, and Experiments. Front. Neural Circuits 11, 108 (2017).

2. Gerstner, W., Lehmann, M., Liakoni, V., Corneil, D. & Brea, J. Eligibility Traces and Plasticity on Behavioral Time Scales: Experimental Support of NeoHebbian Three-Factor Learning Rules. Front. Neural Circuits 12, (2018).

3. Mackintosh, N. J. Blocking of conditioned suppression: Role of the first compound trial. J. Exp. Psychol. Anim. Behav. Process. 1, 335–345 (1975).

4. Rothkopf, C. A. & Ballard, D. H. Credit Assignment in Multiple Goal Embodied Visuomotor Behavior. Front. Psychol. 1, (2010).

5. Crow, T. J. Cortical Synapses and Reinforcement: a Hypothesis. Nature 219, 736–737 (1968).

6. Fremaux, N., Sprekeler, H. & Gerstner, W. Functional Requirements for Reward-Modulated Spike-Timing-Dependent Plasticity. J. Neurosci. 30, 13326–13337 (2010).

7. Izhikevich, E. M. Solving the Distal Reward Problem through Linkage of STDP and Dopamine Signaling. Cereb. Cortex 17, 2443–2452 (2007).

8. Gavornik, J. P., Shuler, M. G. H., Loewenstein, Y., Bear, M. F. & Shouval, H. Z. Learning reward timing in cortex through reward dependent expression of synaptic plasticity. Proc. Natl. Acad. Sci. 106, 6826–6831 (2009).

9. Frémaux, N. & Gerstner, W. Neuromodulated Spike-Timing-Dependent Plasticity, and Theory of Three-Factor Learning Rules. Front. Neural Circuits 9, 85 (2015).

10. Cassenaer, S. & Laurent, G. Conditional modulation of spike-timing-dependent plasticity for olfactory learning. Nature 482, 47–52 (2012).

11. Shindou, T., Shindou, M., Watanabe, S. & Wickens, J. A silent eligibility trace enables dopamine-dependent synaptic plasticity for reinforcement learning in the mouse striatum. Eur. J. Neurosci. 49, 726–736 (2019).

12. Yagishita, S. et al. A critical time window for dopamine actions on the structural plasticity of dendritic spines. Science 345, 1616–1620 (2014).

13. Brzosko, Z., Schultz, W. & Paulsen, O. Retroactive modulation of spike timingdependent plasticity by dopamine. Elife 4, (2015).

14. Brzosko, Z., Zannone, S., Schultz, W., Clopath, C. & Paulsen, O. Sequential neuromodulation of Hebbian plasticity offers mechanism for effective reward-based navigation. Elife 6, (2017).

15. He, K. et al. Distinct Eligibility Traces for LTP and LTD in Cortical Synapses. Neuron 88, 528–538 (2015).

16. Fisher, S. D. et al. Reinforcement determines the timing dependence of corticostriatal synaptic plasticity in vivo. Nat. Commun. 8, 334 (2017).

17. Yoshida, T., Ozawa, K. & Tanaka, S. Sensitivity profile for orientation selectivity in the visual cortex of goggle-reared mice. PLoS One 7, e40630 (2012).

18. Girman, S. V, Sauvé, Y. & Lund, R. D. Receptive field properties of single neurons in rat primary visual cortex. J. Neurophysiol. 82, 301–11 (1999).

19. Nadim, F. & Bucher, D. Neuromodulation of neurons and synapses. Curr. Opin. Neurobiol. 29, 48–56 (2014).

20. Frenkel, M. Y. & Bear, M. F. How Monocular Deprivation Shifts Ocular Dominance in Visual Cortex of Young Mice. Neuron 44, 917–923 (2004).

21. Lehmann, K. & Löwel, S. Age-dependent ocular dominance plasticity in adult mice. PLoS One 3, e3120 (2008).

22. Sato, M. & Stryker, M. P. Distinctive features of adult ocular dominance plasticity. J. Neurosci. 28, 10278–86 (2008).

23. Smith, G. B., Heynen, A. J. & Bear, M. F. Bidirectional synaptic mechanisms of ocular dominance plasticity in visual cortex. Philos. Trans. R. Soc. Lond. B. Biol. Sci. 364, 357–67 (2009).

24. Turrigiano, G. G. & Nelson, S. B. Homeostatic plasticity in the developing nervous system. Nat. Rev. Neurosci. 5, 97–107 (2004).

25. Cang, J., Kalatsky, V. A., Löwel, S. & Stryker, M. P. Optical imaging of the intrinsic signal as a measure of cortical plasticity in the mouse. Vis. Neurosci. 22, 685–91 (2005).

26. Goltstein, P. M., Coffey, E. B. J., Roelfsema, P. R. & Pennartz, C. M. A. In Vivo Two-Photon Ca2+ Imaging Reveals Selective Reward Effects on Stimulus-Specific Assemblies in Mouse Visual Cortex. J. Neurosci. 33, 11540–11555 (2013).

27. Goltstein, P. M., Meijer, G. T. & Pennartz, C. M. Conditioning sharpens the spatial representation of rewarded stimuli in mouse primary visual cortex. Elife 7, (2018).

28. Seitz, A. R., Kim, D. & Watanabe, T. Rewards evoke learning of unconsciously processed visual stimuli in adult humans. Neuron 61, 700–7 (2009).

29. Jurjut, O., Georgieva, P., Busse, L. & Katzner, S. Learning Enhances Sensory Processing in Mouse V1 before Improving Behavior. J. Neurosci. 37, 6460–6474 (2017).

30. Henschke, J. U. et al. Reward Association Enhances Stimulus-Specific Representations in Primary Visual Cortex. Curr. Biol. 30, 1866–1880.e5 (2020).

31. Shuler, M. G. & Bear, M. F. Reward timing in the primary visual cortex. Science 311, 1606–9 (2006).

32. Roelfsema, P. R., van Ooyen, A. & Watanabe, T. Perceptual learning rules based on reinforcers and attention. Trends Cogn. Sci. 14, 64–71 (2010).

33. Roelfsema, P. R. & Holtmaat, A. Control of synaptic plasticity in deep cortical networks. Nat. Rev. Neurosci. 19, 166–180 (2018).

34. Huertas, M. A., Schwettmann, S. E. & Shouval, H. Z. The Role of Multiple Neuromodulators in Reinforcement Learning That Is Based on Competition between Eligibility Traces. Front. Synaptic Neurosci. 8, 37 (2016).

35. Boureau, Y.-L. & Dayan, P. Opponency revisited: competition and cooperation between dopamine and serotonin. Neuropsychopharmacology 36, 74–97 (2011).

36. Matias, S., Lottem, E., Dugué, G. P. & Mainen, Z. F. Activity patterns of serotonin neurons underlying cognitive flexibility. Elife 6, (2017).

37. Clarke, H. F., Dalley, J. W., Crofts, H. S., Robbins, T. W. & Roberts, A. C. Cognitive inflexibility after prefrontal serotonin depletion. Science 304, 878–80 (2004).

38. Kasamatsu, T. & Pettigrew, J. D. Depletion of brain catecholamines: failure of ocular dominance shift after monocular occlusion in kittens. Science 194, 206–9 (1976).

39. Bear, M. F. & Singer, W. Modulation of visual cortical plasticity by acetylcholine and noradrenaline. Nature 320, 172–176 (1986).

40. Bakin, J. S. & Weinberger, N. M. Induction of a physiological memory in the cerebral cortex by stimulation of the nucleus basalis. Proc. Natl. Acad. Sci. 93, 11219–11224 (1996).

41. Kilgard, M. P. Cortical Map Reorganization Enabled by Nucleus Basalis Activity. Science 279, 1714–1718 (1998).

42. Shulz, D. E., Sosnik, R., Ego, V., Haidarliu, S. & Ahissar, E. A neuronal analogue of state-dependent learning. Nature 403, 549–53 (2000).

43. Gu, Q. Neuromodulatory transmitter systems in the cortex and their role in cortical plasticity. Neuroscience 111, 815–35 (2002).

44. Martins, A. R. O. & Froemke, R. C. Coordinated forms of noradrenergic plasticity in the locus coeruleus and primary auditory cortex. Nat. Neurosci. 18, 1483–92 (2015).

45. Huang, S. et al. Pull-Push neuromodulation of LTP and LTD enables bidirectional experience-induced synaptic scaling in visual cortex. Neuron 73, 497–510 (2012).

46. Hong, S. Z., Huang, S., Severin, D. & Kirkwood, A. Pull-push neuromodulation of cortical plasticity enables rapid bi-directional shifts in ocular dominance. Elife 9, (2020).

47. Nakadate, K., Imamura, K. & Watanabe, Y. c-Fos activity mapping reveals differential effects of noradrenaline and serotonin depletion on the regulation of ocular dominance plasticity in rats. Neuroscience 235, 1–9 (2013).

48. Mery, F. A Cost of Long-Term Memory in Drosophila. Science 308, 1148–1148 (2005).

49. Placais, P.-Y. & Preat, T. To Favor Survival Under Food Shortage, the Brain Disables Costly Memory. Science 339, 440–442 (2013).

50. Plaçais, P.-Y. et al. Upregulated energy metabolism in the Drosophila mushroom body is the trigger for long-term memory. Nat. Commun. 8, 15510 (2017).

51. Li, H. L. & van Rossum, M. C. Energy efficient synaptic plasticity. Elife 9, (2020).

